# KaMLs for Predicting Protein p*K*_a_ Values and Ionization States: Are Trees All You Need?

**DOI:** 10.1101/2024.11.09.622800

**Authors:** Mingzhe Shen, Daniel Kortzak, Simon Ambrozak, Shubham Bhatnagar, Ian Buchanan, Ruibin Liu, Jana Shen

## Abstract

Despite its importance in understanding biology and computer-aided drug discovery, the accurate prediction of protein ionization states remains a formidable challenge. Physics-based approaches struggle to capture the small, competing contributions in the complex protein environment, while machine learning (ML) is hampered by scarcity of experimental data. Here we report the development of p*K*_a_ ML (KaML) models based on decision trees and graph attention networks (GAT), exploiting physicochemical understanding and a new experiment p*K*_a_ database (PKAD-3) enriched with highly shifted p*K*_a_’s. KaML-CBtree significantly outperforms the current state of the art in predicting p*K*_a_ values and ionization states across all six titratable amino acids, notably achieving accurate predictions for deprotonated cysteines and lysines – a blind spot in previous models. The superior performance of KaMLs is achieved in part through several innovations, including separate treatment of acid and base, data augmentation using AlphaFold structures, and model pretraining on a theoretical p*K*_a_ database. We also introduce the classification of protonation states as a metric for evaluating p*K*_a_ prediction models. A meta-feature analysis suggests a possible reason for the lightweight tree model to outperform the more complex deep learning GAT. We release an end-to-end p*K*_a_ predictor based on KaML-CBtree and the new PKAD-3 database, which facilitates a variety of applications and provides the foundation for further advances in protein electrostatics research.

**TOC Graphic:** 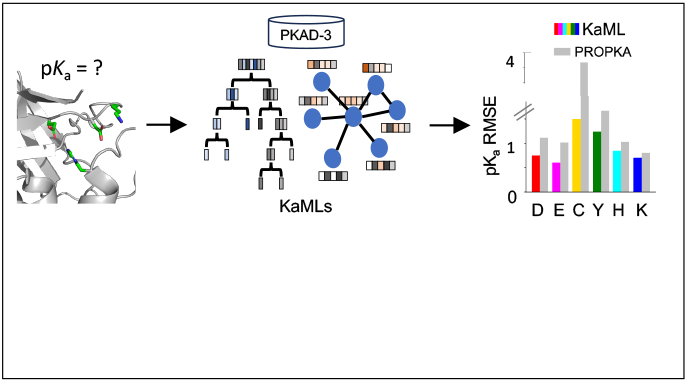

## INTRODUCTION

Ionizable residues in proteins play a variety of roles in biology. For example, enzyme reactions often involve catalytic acid and base, which can donate or abstract a proton,^1^ while pH-dependent ion channels and transporters sense the pH gradient through the protonation or deprotonation of gating residues.^2,3^ Redox processes often involve reactive Cys that is deprotonated or has a high tendency to deprotonate at physiological pH.^4^ Ionizable residues offer unique opportunities for targeted covalent drug discovery. In particular, the deprotonated Cys and Lys residues are nucleophilic, making them valuable targets for covalent inhibitors.^5^ Protein ionization equilibria are characterized by the p*K*_a_ values, which may significantly deviate from the solution (also called model) value. Thus, knowledge of protein p*K*_a_ values is important.

Solution NMR is the method of choice for the determination of site-specific p*K*_a_;^6^ however, it is costly and time consuming. Computational methods offer a potential alternative; however, achieving accuracy and efficiency in p*K*_a_ calculations remains a formidable challenge.^7^ Several physics-based p*K*_a_ prediction approaches have been developed in the past.^7^ A classic approach is based on solving the Poisson-Boltzmann (PB) equation; popular software tools include H++,^8^ DelPhiPKa,^9^ PDB2PQR,^10^ MCCE2,^11^ and PypKa.^12^ One major limitation is the assumption of a uniform protein dielectric constant. In reality, this constant varies from the interior to the surface of the protein.^13^ Significantly faster than PB solvers are empirical methods based on energy functions, e.g., the popular PROPKA program^14,15^ calculates the p*K*_a_ shifts relative to the model values using contributions from desolvation, hydrogen-bonding (h-bond), and charge-charge interactions. Arguably the most accurate and time-consuming p*K*_a_ calculation method^7^ is based on constant pH molecular dynamics (MD) simulations,^16^ e.g., the generalized Born (GB)^17^ or all-atom particle Ewald continuous constant pH MD (CpHMD).^18–20^

In recent years, machine learning (ML) models for p*K*_a_ predictions have emerged as an alternative to physics-based approaches; however, building ML p*K*_a_ predictors is challenging due to the lack of experimental data. Alexov and coworker spearheaded the effort to curate experimental p*K*_a_’s and published the first database PKAD,^21^ which was recently expanded to PKAD-2.^22^ PKAD-2 contains 1,742 entries; however, due to the inclusion of multiple protein data bank (PDB) structures per protein and/or multiple p*K*_a_ measurements per residue, these entries correspond to only 615 unique residues in 113 unique wild-type (WT) or mutant proteins (Table 1). Moreover, the majority of p*K*_a_’s belong to Asp (175), Glu (218), and His (116), while only 20 Cys, 19 Tyr, and 67 Lys are included (Table 1). Furthermore, most of the p*K*_a_’s cluster around the model values and significantly shifted p*K*_a_’s are rare (Figure 1A), making it challenging to train ML models capable of predicting large p*K*_a_ shifts, which are often crucial for biological functions.

**Table 1:** Statistics^*a*^ of p*K*_a_ databases PKAD-2 and PKAD-3.

|  | PKAD-2<sup>b</sup> |  |  | PKAD-3 |  |  | % incr. |  |  |
| --- | --- | --- | --- | --- | --- | --- | --- | --- | --- |
|  | Res | pK<sub>a</sub>'s | PDBs | Res | pK<sub>a</sub>'s | PDBs | Res | pK<sub>a</sub>'s | PDBs |
| Asp | 175 | 214 | 403 | 291 | 330 | 520 | 66.3% | 54.2% | 29.0% |
| Glu | 218 | 258 | 447 | 342 | 382 | 580 | 56.9% | 48.1% | 29.8% |
| His | 116 | 170 | 243 | 155 | 219 | 293 | 33.6% | 28.8% | 20.6% |
| Cys | 20 | 20 | 20 | 57 | 60 | 62 | 185.0% | 200% | 210% |
| Tyr | 19 | 20 | 22 | 38 | 39 | 41 | 100.0% | 95.0% | 86.4% |
| Lys | 67 | 81 | 151 | 109 | 137 | 216 | 62.7% | 69.1% | 43.0% |
| Total | 615 | 763 |  | 992 | 1167 |  | 61.3% | 52.9% |  |
| Entries |  |  | 1286 |  |  | 1712 |  |  | 33.1% |
| Proteins |  |  | 113 |  |  | 247 |  |  | 118.6% |

<sup>a</sup>The number of unique residues, the number of measured pK<sub>a</sub>'s, the number of PDB entries (X-ray structures), and the number of unique wild-type or mutant proteins. The number of PDBs includes several modeled structures for mutant proteins. <sup>b</sup>A cleaned version of PKAD-2,<sup>22</sup> where pK<sub>a</sub>'s given in ranges were removed and errors were corrected (see Methods).

**Figure 1:**
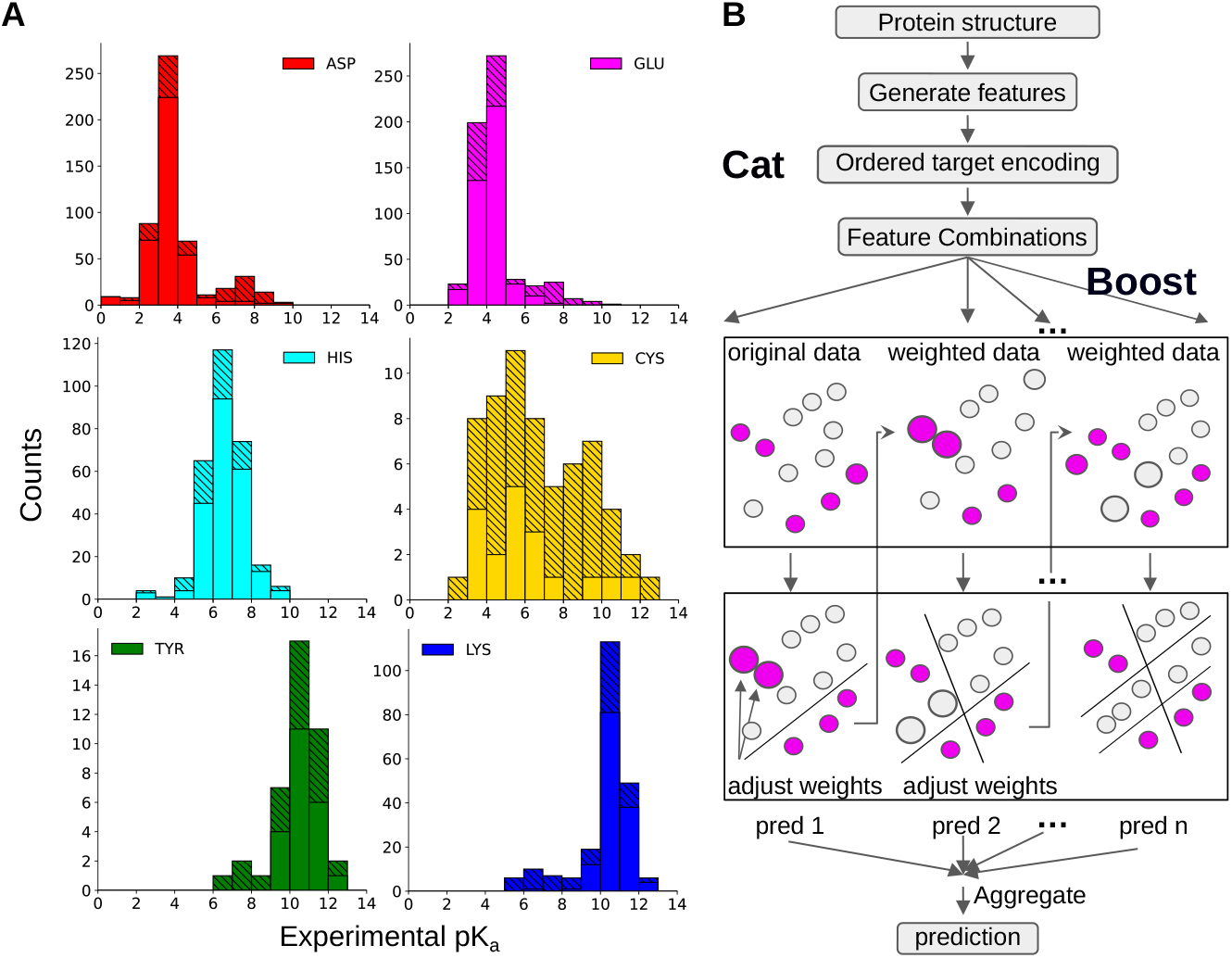
Overview of the p*K*_a_ dataset and illustration of KaML-CBtree. **A**. Histograms of the experimental p*K*_a_ values in PKAD-3 database. Solid bars represent the data in the cleaned PKAD-2 database and striped bars represent the data added in this work. **B**. The Catboost algorithm. CatBoost handles categorical features using ordered target statistics to capture the relationship with the target variable. Gradient boosting algorithms sequentially build an ensemble of decision trees, where each tree corrects the errors of the previous ones by adjusting the weights for data points with large prediction errors. The final aggregated prediction is the sum of the individual predictions of all trees.

To tackle data scarcity, Reis, Machuqueiro et al. developed pKPDB,^23^ a theoretical p*K*_a_ database comprised of 12 million p*K*_a_’s of six titratable amino acids calculated using PypKa,^12^ a Python API for the fast PB solver DelPhi v5.^24^ Based on the p*K*_a_ shifts from pKPDB, Reis, Machuqueiro, Clevert et al. trained PKAI+ (a multilayer perceptron),^25^ which achieved root-mean-square error (RMSE) of 0.98 on a subset of 750 p*K*_a_’s from the PKAD database.^21^ Using pH replica-exchange^26^ GBNeck2-CpHMD titration simulations,^27,28^ Huang and coworkers created a theoretical database PHMD549^29,30^ comprising 27k p*K*_a_’s of Asp, Glu, His, and Lys from 549 proteins in the latest version.

Recently, ML models trained on experimental p*K*_a_ data have also been reported. Based on PKAD^21^ and 23 additional mutant p*K*_a_’s, Chen, Lee, Damjanovic et al. trained decision tree models using 12 features.^31^ After correcting for training/test data leakage, the RMSE of the best model XGBoost is above 1 (Damjanovic, Protein Electrostatics Conference 2023, Genoa, Italy). Based on PKAD,^21^ Yang, Luo and coworker^32^ trained XGBoost tree model using atom-based and distance features, which gave RMSE of 1.0 for Asp, Glu, His, and Lys when evaluated on 20% of the unseen PKAD data. Isayev and coworker developed (shallow) neural network p*K*_a_ predictors^33^ using an atomic environment vector embedding model called ANI-2x,^34^ which is an extension of the original ANI model^35^ by including three more chemical element types. The ANI-2x^34^ (or ANI)^35^ model is trained to reproduce quantum mechanical potential energy surfaces of small molecules. Given a protein structure, the atomic environment vector represents the bonded and non-bonded interactions within the default ANI cutoff distance of 5.2 Å.^33^ Trained on a subset of PKAD^21^ for individual residue types, the ANI-2x p*K*_a_ predictors^33^ achieved RMSEs of 0.59 for Glu, 0.94 for Asp, and 0.98 for His in one holdout test.

Building on the aforementioned advances, this work addresses several key challenges in p*K*_a_ predictions. First, we developed a new database PKAD-3, which expands on and refines PKAD-2.^22^ Next, we developed p***K***_**a**_ **M**achine **L**earning models (KaMLs), using the CatBoost decision tree (CBtree) model and to our best knowledge the first graph attention network (GAT),^36^ for protein p*K*_a_ predictions. KaML-CBtree and KaML-GAT achieve state-of-the-art performance through several innovations, such as the separate models for acidic and basic residues; incorporation of features representative of p*K*_a_ determinants; data augmentation with AlphaFold2 (AF2) structures;^37^ and model pretraining on a theoretical p*K*_a_ dataset. Note that AF2 structures have been used to perform proteomewide ML pKa predictions^31^ and most recently to extend the theoretical PypKa database.^38^

Our work eliminated a major data leakage issue in the previous ML model training,^31–33^ whereby residues with different identification numbers (IDs) in different PDB files were treated as distinct residues, while in fact they are the same. Furthermore, we evaluated the models using 20 holdout sets to provide robust performance statistics, whereas the previous ML models were evaluated using only a single holdout set.^25,30–32^ To enhance practical relevance, we extended our model evaluation to include metrics for protonation-state predictions. We then benchmarked the KaMLs against the baseline models (null model and PROPKA3^15^) and the PB solver (PypKa^12^) as well as two different types of ML models (DeepKa^30^ and ANI-2X^33^). Finally, using meta-feature analysis, we rationalized the superior performance of the light-weight KaML-CBtree over the more complex KaML-GAT.

## Results and Discussion

### Development of PKAD-3 and analysis of experimental p*K*_a_ values

#### Developing an expanded, high-quality experimental p*K*_a_ database PKAD-3

An extensive and high-quality dataset is of utmost importance for training and testing ML models. Following manual verification and error correction of entries in PKAD-2,^22^ a literature search was conducted for additional experimental p*K*_a_ values. We refer to the expanded database as PKAD-3, in tribute to the pioneering efforts of the Alexov group.^21,22^ PKAD-3 contains 1167 p*K*_a_’s of 992 unique residues in 247 proteins (WT or mutant), representing increases of 53%, 61%, and 119% over the corresponding numbers (763 p*K*_a_’s of 615 residues in 113 proteins) in the cleaned PKAD-2 (Table 1). Note that since some residues and p*K*_a_’s are associated with more than one PDB structure, PKAD-2 contains a total of 1286 PDB entries, which was mistakenly cited as the number of p*K*_a_’s in the previous publications.^22,25,31–33^ To facilitate community efforts in developing p*K*_a_ prediction tools, we implemented PKAD-3 as a freely searchable and downloadable web database (http://database.computchem.org/pkad3).

#### Analyzing experimental p*K*_a_ distributions of six titratable amino acids

Compared to PKAD-2, there is a substantial increase in the number of (unique) residues with experimental p*K*_a_’s for all six titratable amino acids, particularly Cys and Tyr (Table 1 and Figure 1A). The number of Cys and Tyr increases by 185% (from 20 to 57) and 100% (from 19 to 38), respectively. The number of Asp, Glu, and Lys increases by 58–67%, while the number of His shows a modest increase of 34%. As in PKAD-2, the number of residues and experimental p*K*_a_’s for Glu, Asp, His, Lys, Cys, and Tyr follows a descending order (Table 1).

Before training ML models, it is instructive to examine the distributions of target data (Figure 1A). The p*K*_a_ distributions of individual amino acids exhibit distinct patterns. His p*K*_a_’s display a nearly Gaussian distribution centered around the model p*K*_a_ of 6.5, demonstrating that His protonated, deprotonated, or titrating at physiological pH. This also suggests that it may be necessary to allow His titration in MD simulations via constant pH techniques.^16^

Cys p*K*_a_’s exhibit a distinctive bimodal distribution with two peaks of similar heights. Surprisingly, the major peak is near p*K*_a_ 5—6, indicating deprotonation at physiological pH. Deprotonated Cys or those with p*K*_a_’s near physiological pH are referred to as hyperreactive or reactive, as they are prone to chemical modifications and play important roles in catalysis and redox chemistry.^4,17,39^ Thus, the relative abundance of the low Cys p*K*_a_’s may be an experimental study bias due to the biological significance. The secondary peak of Cys p*K*_a_’s is located at 9–10, indicating protonation at physiological pH. The unique Cys p*K*_a_ distribution, combined with the small dataset, presents a challenge for ML models. The prevalence of deprotonated Cys suggests a potential source of inaccuracy in MD simulations, since the protonated form is the default state.

The p*K*_a_ distributions of Asp, Glu, Tyr, and Lys are bimodal, with a dominant peak near the model p*K*_a_ and a minor peak at a higher (Asp and Glu) or lower (Tyr and Lys) p*K*_a_’s. The p*K*_a_ ranges of Asp and Glu extend to approximately 9, demonstrating that Asp and Glu can titrate or even protonate under physiological conditions. Most of these large p*K*_a_’s were obtained by the García-Moreno lab through NMR or denaturation experiments of engineered mutants of staphylococcal nuclease (SNase), whereby a buried hydrophobic residue is replaced with a titratable one.^40–42^

The mutant SNase experiments also contributed Lys p*K*_a_’s that extend down to values as low as 5,^43,44^ indicating that Lys can titrate or deprotonate under physiological conditions. In addition to producing large p*K*_a_ shifts, some of the mutated residues in SNase show normal (i.e. near model) p*K*_a_’s despite being buried in the interior.^45^ These normal and significantly shifted p*K*_a_’s are collectively termed anomalous p*K*_a_’s; they provide crucial training data for ML models to generalize across diverse protein environments and capture rare but functionally relevant p*K*_a_’s. Note, anomalous p*K*_a_’s are most challenging to predict accurately using physics-based approaches^7^ and may require accurate description of ionization-induced conformational changes.^45^

Similarly to Lys, but to a smaller extent, the p*K*_a_ distribution of Tyr displays low values of 6–9, indicating that some Tyr can titrate or even deprotonate at physiological pH. Due to the extremely small dataset, it is challenging to train ML models that can predict deprotonated Tyr.

### Training and evaluating KaML-trees

#### Feature engineering and visualization

Due to the small dataset (1,167 p*K*_a_’s in PKAD-3), we first turned to the “shallow learning” tree models, which recursively divide features into subsets, forming a tree-like structure of decisions. We adapted our recent tree features for predicting Cys ligandabilities,^46^ which include three types of numerical features that represent the physical determinants of p*K*_a_ shifts:^15,41,47^ solvent accessibility, potential hydrogen bonding, and electrostatic interactions. In addition, categorical features that describe residue type, net charge, and secondary structures in proximity to the titratable residue were included (see complete list in SI Methods and Table S1).

To test whether tree features exhibit clusters that are correlated with protonation states, we used the stochastic neighbor embedding (t-SNE) algorithm,^48,49^ which calculates pairwise similarities between data points in the high-dimensional space and maps them to a lower-dimensional space while preserving the neighbor identities.^48,49^ His p*K*_a_’s are clustered near 7, and therefore we used their features as a stringent test case. Using t-SNE, the 37 numerical features are mapped on two dimensions, where each point is colored by the (target) protonation state (Figure 2 for pH 7 and SI Figure S2 for pH 7.5). The plot displays several clusters, each predominantly characterized by a single color, suggesting that the features can effectively discriminate between distinct protonation states.

**Figure 2:**
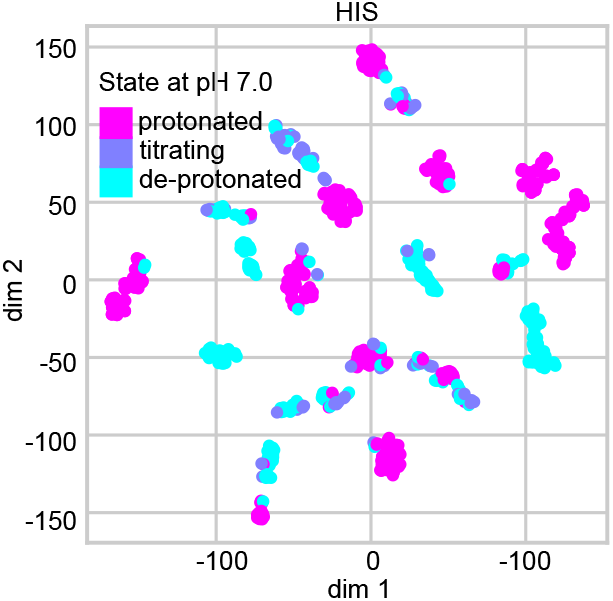
t-SNE visualization of the histidine features. The 37 numerical features of His were extracted from the dataset with the AF2 augmentation and mapped on two dimension using t-SNE. The color code indicates the protonation state at pH 7 based on the label (experimental) p*K*_a_: deprotonated (p*K*_a_ *>* 7.48); protonated (p*K*_a_ *<* 6.52); and titrating (6.52 ≤ p*K*_a_ ≤ 7.48).

#### Data splitting, training and evaluating the acid and base KaML-trees

We separated the data into acidic (Asp, Glu, Cys, Tyr) and basic (His and Lys) residues due to the distinct mechanisms of p*K*_a_ shifts and trained separate models for them (see the later discussion). This resulted in 728 acidic residues (811 p*K*_a_’s and 1203 PDB entries) and 264 basic residues (356 p*K*_a_’s and 509 PDB entries). The data was randomly split into 90% for training and 10% for holdout test (SI Figure S1). To prevent data leakage, data splitting was based on unique residues (UniProt ID + UniProt + resid + mutant + conformational states) and stratification was applied based on the p*K*_a_’s (see Methods for details). Data splitting based on protein sequences is sensible but was not pursued for the following reasons. First, protein p*K*_a_’s are determined by the local electrostatic environment, i.e., two residues in a similar sequence environment may not necessarily have similar p*K*_a_’s due to a nearby mutation or a large conformational change. Another case involves engineered titratable residues such as those in SNase made by the Garcia-Moreno lab,^40–42^ which can have drastically different p*K*_a_’s despite having a similar background protein sequence. Second, due to the limited training dataset size, sequence-based data splitting would reduce the number of samples too severely to be practical. Finally, a large number of holdout tests with different random splits can minimize potential data leakage between training and test.

We first trained five types of tree models: Random forest (RF), Extra trees (ET), gradient boosting (GB), extreme gradient boosting (XGB) and Catboost (CB). Each model was trained on each of the 20 different train/test splits and 10 fold cross-validation (CV) was used during training. Mean square error (MSE) of p*K*_a_’s was used as training loss. The average and standard deviation of PCC, RMSE, and absolute maximum error (MAXE) from 20 holdout tests were used as evaluation metrics.

We first compared the metrics of the different tree models (SI Table S2). Considering all residues, the PCCs of all five tree models are nearly identical (0.94); however, the CBtree yields the lowest RMSE (0.77) and MAXE (3.47), followed by ETtree, which yields RMSE of 0.80 and MAXE of 3.70. Considering acid and base separately, the acid CBtree remains the best, with the RMSE of 0.77 and MAXE of 3.25, as compared to 0.82 and 3.61 with the ETtree, although the RMSE and MAXE of the base CBtree (0.76 and 2.60) are slightly higher than those of the base ETtree (0.74 and 2.46).

For practical applications, the exact p*K*_a_ is often less relevant than the protonation state at a specific pH, e.g., the physiological cytosolic pH of 7.1. Thus, we evaluated the model’s performance of correctly predicting protonation states by discretizing the predicted p*K*_a_’s in three classes based on the protonation probability (Prob) at pH 7: protonated (Prob *>* 0.75 or p*K*_a_*<* 6.52), deprotonated (Prob *<* 0.25 or p*K*_a_*>* 7.48), or titrating (0.25 ≤ Prob ≤ 0.75 or 6.52 ≤ p*K*_a_ ≤ 7.48). This discretization step effectively transforms the regression problem into a classification task, and consequently, the models can be evaluated by the class precision (Pre) and recall (Rec).

Here we concern ourselves with the protonated and deprotonated classes and defer the discussion of the titrating class to future work. Rec informs the percentage of protonated (deprotonated) residues identified, while Pre informs the correct percentage of the identified protonated (deprotonated) residues. Since incorrectly predicting protonated as deprotonated or vice versa is the most consequential error, we evaluated the percentage of these “misclassified” instances and referred to it as the critical error rate (CER).

Considering Pre/Rec for protonated and deprotonated classes of all residues (SI Table S3), the CBtree again emerged as the best performing tree model, yielding Pre/Rec of 0.97*/*0.93 for the protonated class and 0.94*/*0.94 for deprotonated class (SI Table S3). The CER of CBtree is also the lowest (46/2635). Considering Pre/Rec and CER of the acid and base models separately (Table 2), the CER of the base CBtree is nearly three times lower than ETtree, although the acid CBtree has slightly lower Rec and higher CER than the acid ETtree. Since CBtree has overall the best performance, we will drop the other tree models in the remainder of the discussion.

**Table 2:**
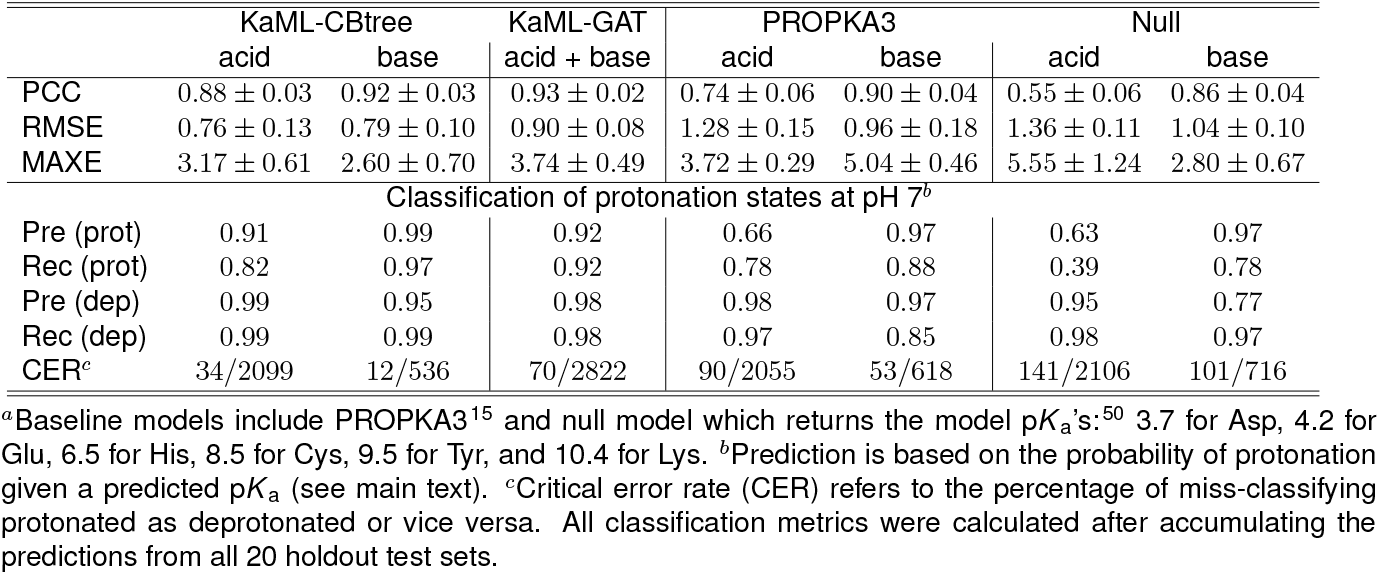
Performance metrics of KaML-CBtree for acid and base p*K*_a_ and protonation state predictions in comparison to the baseline models^*a*^.

#### KaML-CBtree outperforms PROPKA3 in p*K*_a_ prediction accuracy across all titratable amino acids

The CBtree significantly outperforms the baseline models, PROPKA3^15^ and null model, in p*K*_a_ predictions. The CBtree yields PCCs of 0.88/0.92, RMSEs of 0.76/0.79, and MAXEs of 3.17/2.60 for acid / base residues, compared to PROPKA3 PCCs of 0.74/0.90, RMSEs of 1.28/0.95, and MAXEs of 3.72/5.04 (Table 2). Importantly, the differences between CBtree and PROPKA3 far exceed the statistical uncertainties of the model evaluations.

The amino acid-specific PCC is a stringent evaluation metric due to the narrow p*K*_a_ range for individual amino acids. The CBtree yields PCCs of 0.86 for Asp, 0.84 for Glu, 0.51 for His, 0.61 for Cys, and 0.80 for Lys, which are higher than PROPKA3’s PCCs of 0.64 for Asp, 0.69 for Glu, 0.45 for His, 0.12 for Cys, and 0.75 for Lys (Figure 3A). The most notable improvement is for Cys, as PROPKA3 predictions do not offer a statistically meaningful correlation with experiment, which is likely due to the extremely small number of experimental Cys p*K*_a_’s used in fitting the PROPKA3 model.^15^

**Figure 3:**
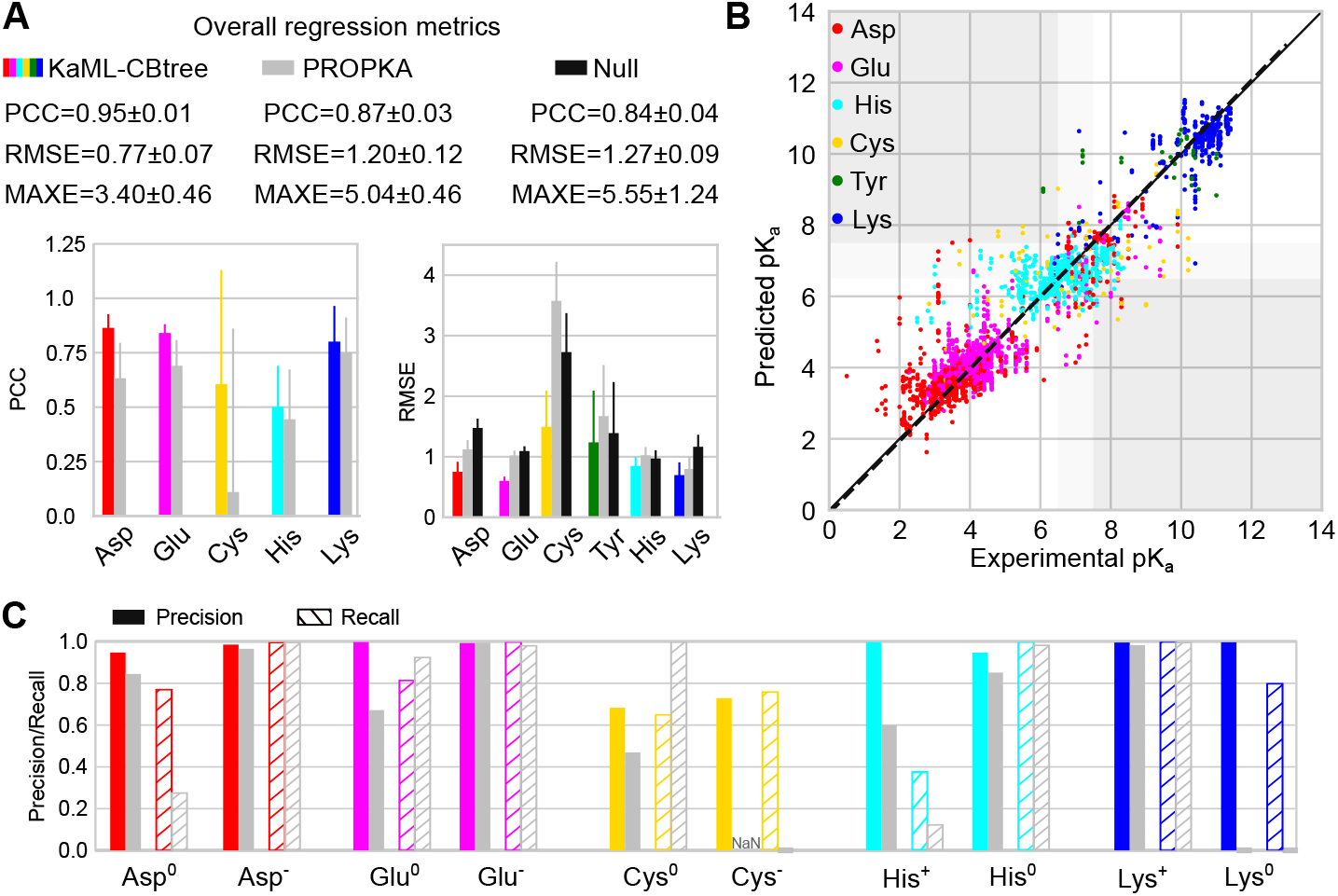
Evaluation of KaML-CBtree for predicting p*K*_a_’s and protonation states of individual titratable amino acids. **A**. Top. Overall performance metrics of KaML-CBtree (color-coded), PROPKA3 (grey), and null model (black). Bottom. PCC and RMSE for individual amino acids. The mean and SD of 20 holdout tests are given. **B**. Experimental vs. predicted p*K*_a_’s from all 20 test sets (see SI Figure S3 for plots of individual test sets). The solid line is the identity and the dotted line is a linear fit. Data points are color-coded by amino acid. The white regions indicate correct protonation state classifications. The dark gray regions highlight the critical errors. The light gray regions indicate titrating p*K*_a_’s which were excluded from the classification analysis. **C**. The precision (solid) and recall (with stripes) of classifying protonated and deprotonated amino acids at pH 7 using KaML-tree (color-coded) and PROPKA3 (grey). ^15^ PROPKA3 fails to predict any Cys^−^; thus precision is undefined (NaN). Classification analysis for Tyr is not shown due to the extremely small test sets.

Comparing the CBtree PCCs across different amino acids, Asp, Glu, and Lys show the highest PCCs around 0.8, whereas the PCCs for His and Cys are much lower, 0.51 and 0.61, respectively. The low PCC for His may be attributed to its narrow p*K*_a_ distribution (Figure 1A), in which case even small errors can decrease the PCC.

Compared to PROPKA3 and the null model, the RMSE of CBtree is lower for all six amino acids (Figure 3A and B). The CBtree yields RMSEs of 0.75 for Asp, 0.60 for Glu, 0.85 for His, 1.50 for Cys, 1.24 for Tyr, and 0.70 for Lys, which are significantly lower than PROPKA3’s RMSEs of 1.12 for Asp, 1.02 for Glu, 1.03 for His, 3.58 for Cys, 1.67 for Tyr, and 0.80 for Lys. The largest RMSE reduction is for Cys, 2.1. In contrast, the smallest RMSE reduction is for His, only 0.18, which is due to the concentration of experimental p*K*_a_ values around the model value (Figure 1A). This is supported by the RMSE of the null model being 0.06 lower than PROPKA3 (Figure 3A).

Comparing the CBtree’s RMSEs across different amino acids, Glu shows the smallest RMSE of 0.60, which approaches the experimental p*K*_a_ error of 0.5 units.^6^ This level of accuracy likely stems from the extensive training dataset (580 p*K*_a_’s of 342 residues; Table 1). The largest RMSE is for Cys (1.50), followed by Tyr (1.24; Figure 3A). This trend is the same for both the CBtree and PROPKA3, reflecting the small size of training (for CBtree) or fitting (for PROKA3) dataset.

#### KaML-CBtree outperforms PROPKA3 in protonation state classification across all six amino acids

Considering acid and base separately, CBtree yields higher Pre and Rec of both protonated and deprotonated states than PROPKA3 and null model in (Table 2). The most dramatic improvement is in the reduction of CER. CBtree’s CER in predicting acid protonation states is 34/2099, which is nearly three times lower than PROPKA3 (90/2055) and more than four times lower than null model (141/2106). CBtree’s CER in predicting base protonation states is 12/536, which is more than four times lower than PROPKA3 (53/618) and more than ten times lower than null model (101/716). For acids, the drastic reduction in CER can be largely attributed to the more precise prediction of protonated acids (Pre of 0.91 for CBtree vs. 0.66 for PROPKA3 and 0.63 for null model). For base, the reduction in CER can be attributed to the higher recall of both protonated and deprotonated base residues.

#### Dramatic improvement in predicting Asp^0^, Glu^0^, Cys^−^, and Lys^0^

We compare CB-tree’s protonation state classification for individual amino acids with PROPKA3 (Figure 3C and SI Table S4 and S5). Null model is not evaluated because it predicts only one state. We first consider acids, Asp, Glu, and Cys. Since Asp^−^ and Glu^−^ are dominant, both CBtree and PROPKA have high Pre/Rec; however, the CBtree has a better performance for predicting Asp^0^ and Glu^0^. In particular, CBtree’s Rec for Asp^0^ (0.77) exceeds PROPKA3’s (0.28) by nearly 0.5, and CBtree’s Pre for Glu^0^ (1.0) exceeds PROPKA3’s (0.67) by 0.33. Significantly, CBtree’s Pre/Rec for Cys^−^ is 0.73/0.76, while PROPKA3 fails to identify any Cys^−^, instead assigning Rec of 1 for Cys^0^.

For base amino acids (His and Lys), CBtree shows the most dramatic improvement in predicting Lys^0^, achieving Pre/Rec of 1.0/0.8, while PROPKA3 fails to identify any. His protonation state at pH 7 is most challenging to predict, due to most p*K*_a_’s concentrating around 6.5. While both models have high Pre/Rec for His^0^, the CBtree predicts His^+^ with Pre/Rec of 1.0/0.37, compared to PROPKA3’s 0.60/0.12. Note, the recall appears very low; however, due to class imbalance the recall from random guess is only 0.24. Finally, CBtree’s CERs are 2.4–6 times lower than PROPKA3’s across five titratable amino acids: 13/31 for Asp, 5/21 for Glu, 11/47 for His, 13/35 for Cys, and 1/6 for Lys.

#### Why does training separate acid and base KaML-CBtrees boost performance?

To understand the necessity of training separate tree models for acid and base residues, we examine the impacts of tree features using the SHapley Additive exPlanations (SHAP)^51^ plots for the unseen tests (Figure 4). As expected, the model p*K*_a_ makes the largest impact on acid and base p*K*_a_ predictions. However, the next-largest contributing features affect the acid and base p*K*_a_ predictions differently. In acid p*K*_a_ predictions, a high buried ratio shifts the SHAP value and predicted p*K*_a_ up, increasing the probability of protonation. A similar impact is made by increasing the number of polar sidechains within 10 Å around the residue of interest (n polar10). Similar to buried ratio and n polar10, the number of polar sidechains within 15 Å (n polar15) is the second largest contributor in base p*K*_a_ predictions; however, increasing its value shifts the SHAP value and predicted p*K*_a_ down, increasing the probability of deprotonation.

**Figure 4:**
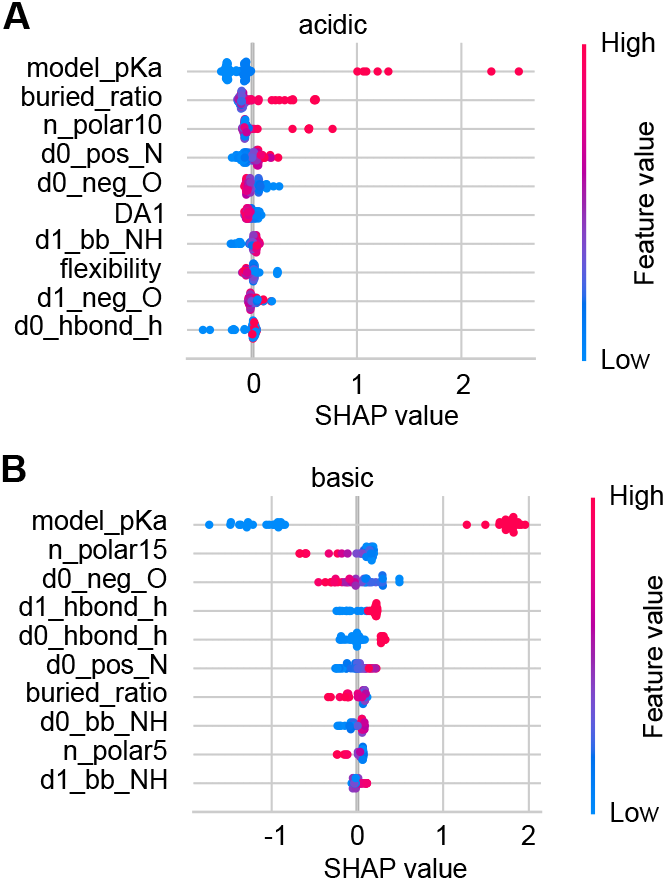
Impacts of features on the p*K*_a_ predictions of acid and base residues. SHAP value plots for the acid (**A**) and base (**B**) p*K*_a_ predictions. The top ten features with the largest average contributions are shown. Each data point is an instance in the test data. Feature values are colored from blue (low) to red (high). The position along the SHAP value axis shows how a feature value shifts the model output. An explanation of all features is given in SI Table S1.

The impacts of opposite signs are also made by features that represent attractive electrostatic interactions. In acid p*K*_a_ predictions, decreasing the distance to the nearest His sidechain nitrogen (d0 hbond h) downshifts the SHAP value and p*K*_a_, increasing the probability of deprotonation. In base p*K*_a_ predictions, decreasing the distance to the nearest negatively charged oxygen (d0 neg O) upshifts the SHAP value and p*K*_a_, increasing the probability of protonation.

The above analysis demonstrates that although important features are shared between acid and base p*K*_a_ predictions, the impacts on p*K*_a_ predictions are in opposite directions. This explains why training separate acid and base KaML-trees offers a significant performance boost.

### Training and evaluating KaML-GAT

#### Building KaML-GATs, data augmentation and model pretraining

In an attempt to further improve the p*K*_a_ and protonation state predictions, we turned to GAT,^36^ an improved version of convolutional graph neural network (GNN). A GNN is designed to process graphstructured data such as protein structures; through message passing steps, the information stored in nodes and edges (node and edge embeddings) can be exchanged with their neighbors. A GAT allows attention to be added based on the neighboring nodes’ features, allowing more weights to be applied to important features. The KaML-GAT architecture and workflow are illustrated in Figure 5A.

**Figure 5:**
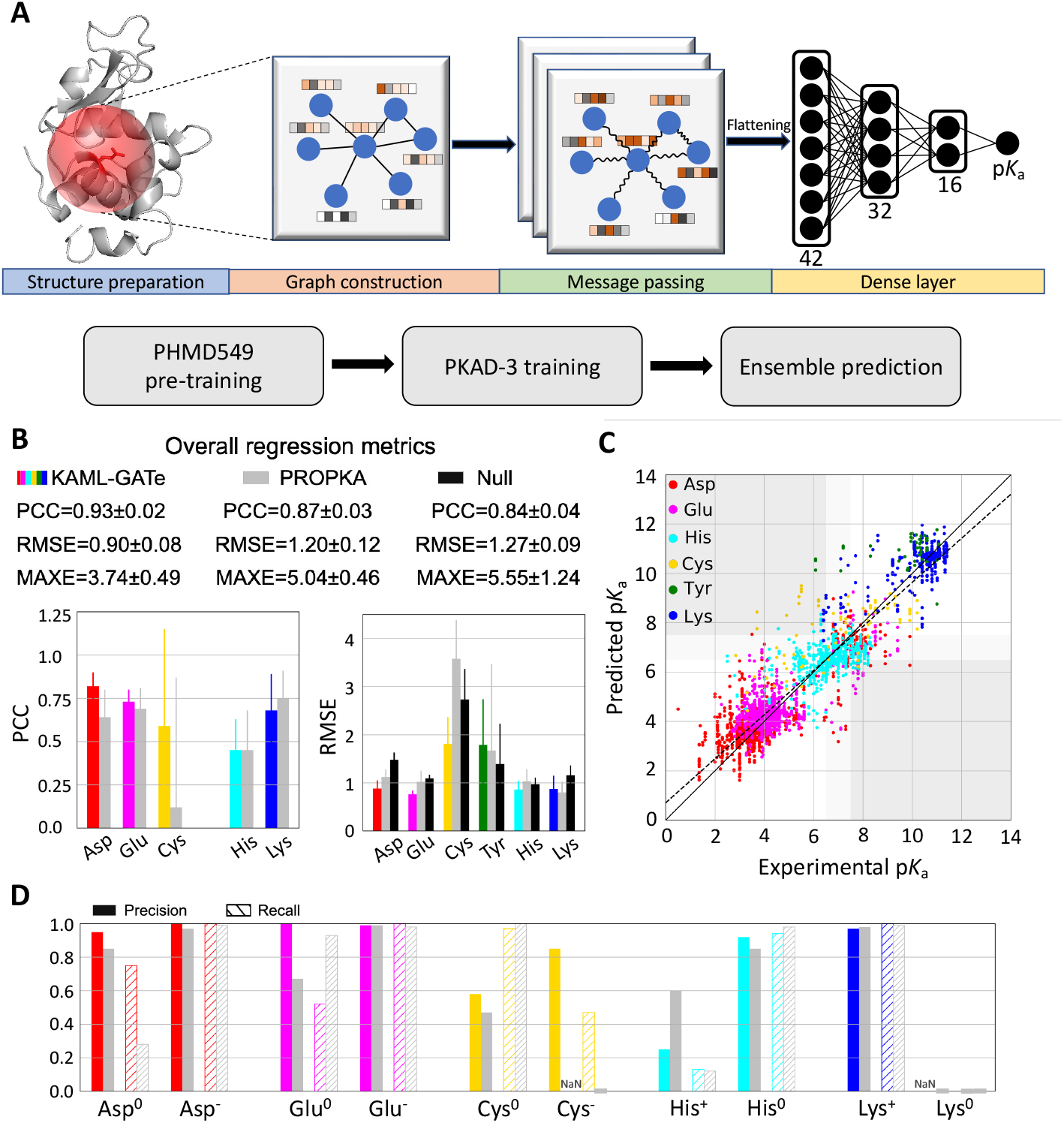
Schematic of KaML-GAT and its performance of predicting p*K*_a_’s and protonation states for five amino acids. **A**. Schematic of KaML-GAT architecture (top) and workflow (bottom). **B**. Overall (top) and amino acid specific PCC and RMSE for p*K*_a_ predictions by KaML-GAT (colored), PROPKA3 (grey), ^15^ and null model (black). **C**. The predicted vs. experimental p*K*_a_’s from 20 unseen tests. The plots for individual tests are given in SI Figure S6. **D**. Precision (solid) and recall (strips) in predicting protonation states at pH 7 for individual amino acids by KaML-GAT and PROPKA3.

For computational efficiency, the protein structure was truncated as a 10-Å sphere around the most relevant atoms of the titratable sidechain, e.g., OD1/OD2 of Asp, NZ of lysine (see Methods for details). Similar to the truncated cubic box (20 Å edges) used in the DeepKa CNN models,^29,30^ this sphere captures the local hydrophobic environment and electrostatic as well as hydrogen bond interactions that may impact the p*K*_a_ values. The 10-Å cutoff reflects the approximate range of electrostatics relevant for p*K*_a_’s, as was used in PROKA3 for Coulomb calculations.^15^ The Graphein package^52^ was used for building the graph. Each node (atom) is represented by a 42-digit vector embedding, similar to the CNN channels for Cys ligandability predictions.^46^ A 24-digit one-hot encoding for the atom types was added (SI Table S6). The message passing is enabled through 3 one-head convolutional layers with 42 channels. Following a global average pooling layer, two hidden layers containing 32 and 16 neurons with a dropout rate of 0.2 are employed before a final layer that makes a p*K*_a_ prediction.

Deep learning requires a large amount of data. To address data shortage, we implemented a two-pronged approach: data augmentation to artificially expand the dataset and model pretraining by leveraging a larger, related dataset. To augment data, we added (based on feature calculations at most 10)

AlphaFold2 (AF2) structure models^37^ for residues with an absolute p*K*_a_ shift greater than 2 (see Methods and SI Figure S4). As target labels, we used p*K*_a_ shifts (reduction in the average test RMSE by 0.3 compared to using p*K*_a_’s as labels, data not shown). Unlike KaML-trees, a single GAT outperformed separate acid and base models, yielding lower test RMSE. For model pretraining, we employed PHMD549, which contains GBNeck2-CpHMD calculated p*K*_a_’s of Asp, Glu, His, and Lys.^30^ After removing the residues in PKAD-3, this dataset contains 26,252 p*K*_a_’s of 25,912 residues in 535 proteins. Data splitting (9:1 ratio for train:test) and holdout test sets are identical to those used for KaML-trees (SI Figure S1). In model pretraining, 10% of data was reserved for evaluation, i.e., model selection and hyperparameter tuning. The pretrained model yields an RMSE of 0.79 for the validation data (SI Figure S5).

To examine the effect of data augmentation (DA) and pretraining (PT) on model performance, we trained additional 3 GATs without DA/PT, with DA only, or with PA only. The results from one train/test split suggest that DA and PA have a synergistic effect on the model performance. Using either DA or PA, the RMSE is increased by about 0.1 while the PCC is decreased by about 0.01, compared to the result with both DA/PT (SI Table S9).

#### The fine-tuned ensemble KaML-GAT out-performs PROPKA3 and null model in overall metrics

Following pretraining on PHMD549, the GAT was fine-tuned by training on the AF2 augmented PKAD-3. holdout tests with 0, 1, or 2 frozen GAT layers demonstrated that releasing all layers gave the lowest overall RMSE and the highest overall PCC (SI Table S7). We also tested the idea of aggregating multiple “weak predictors” to enhance performance.^53^ Specifically, for each training set, 10 models were trained using 10 different training:validation (9:1 ratio) splits, and this process was repeated for each train:holdout data split (20 times total). The ensemble average RMSE decreases and plateaus as the model number reaches 8 (SI Figure S7). Thus, we further evaluated the 10-model based ensemble GAT. For simplicity, we drop the word ensemble in the remainder of the discussion.

The GAT outperforms PROPKA3 and null model in the overall PCC, RMSE, and MAXE (Table 2). The largest improvement is in RMSE/MAXE. The GAT yields RMSE/MAXE of 0.90/3.74, compared to 1.20/5.04 by PROPKA3 and 1.27/5.55 by null model. In the classification of the deprotonation states at pH 7, the GAT also outperforms PROPKA3 and null model. The largest improvement is in CER. Out of 2822 instances, GAT’s CER is 70, which is twice and three times lower than PROPKA (143) and null model (242), respectively.

#### KaML-GAT outperforms PROPKA3 for Asp, Glu, and Cys but not for His, Lys and Tyr

The GAT predicts more accurately p*K*_a_’s for Asp, Glu, and Cys (Figure 5B and D and SI Table S8). The GAT’s PCC/RMSE are 0.82/0.88 (Asp), 0.73/0.76 (Glu), and 0.59/1.81 (Cys), compared to PROPKA3’s 0.64/1.12 (Asp), 0.69/1.02 (Glu), and 0.12/3.58 (Cys). The GAT yields significantly higher Rec (0.75) for Asp^0^, compared to 0.28 with PROPKA3 (Figure 5D). As to Glu^0^, the GAT gives a higher Pre (1.0), compared to 0.67 with PROPKA3; however, the GAT’s Rec (0.52) is lower than that of PROPKA3 (0.93). In predicting Cys^−^, the GAT yields Pre/Rec of 0.85/0.47, whereas PROPKA3 fails to predict Cys^−^.

Surprisingly, the GAT has similar performances as PROPKA3 for His, Lys and Tyr (Figure 5B and D). Although the GAT gives a slightly lower RMSE for His p*K*_a_’s, the Pre of predicting His^+^ is only 0.25, lower than PROKA3’s 0.60. Like PROPKA3, the GAT fails to predict Lys^0^, and the RMSE for Tyr p*K*_a_’s is slightly higher.

### Comparison between KaML-tree and KaML-GAT

#### Performance comparison between KaML-CBtree and KaML-GAT

Both the overall and amino acid specific metrics demonstrate that the CBtree outperforms the GAT (Table 2, Figure 3, Figure 5, and Table 3). Except for His, which shows similar RMSEs between the two models, the CBtree gives lower RMSEs for individual amino acids. Importantly, the CBtree’s CERs for all but Asp are substantially reduced compared to the GAT (Table 3).

**Table 3:** Performance comparison of KaML-CBtree, KaML-GAT, PB, and alternative ML models^*a*^.

|  | PypKa |  | DeepKa |  | ANI-2X<sup>b</sup> |  | KaML-CBtree |  | KaML-GAT |  |
| --- | --- | --- | --- | --- | --- | --- | --- | --- | --- | --- |
|  | RMSE | CER | RMSE | CER | RMSE | CER | RMSE | CER | RMSE | CER |
| Asp | 1.61 ± 0.28 | 56/917 | 1.23 ± 0.35 | 48/937 | 1.20 ± 0.14 | 52/929 | 0.75 ± 0.17 | 13/916 | 0.88 ± 0.17 | 5/901 |
| Glu | 0.86 ± 0.12 | 25/1039 | 0.84 ± 0.25 | 9/1068 | 0.81 ± 0.12 | 40/1075 | 0.60 ± 0.07 | 5/1076 | 0.76 ± 0.08 | 12/1053 |
| His | 1.13 ± 0.51 | 8/257 | 1.10 ± 0.49 | 12/248 | 0.52 ± 0.16 | 3/298 | 0.85 ± 0.14 | 11/209 | 0.86 ± 0.18 | 26/203 |
| Cys | 3.15 ± 0.97 | 21/56 | n/a | n/a | n/a | n/a | 1.50 ± 0.60 | 13/68 | 1.81 ± 0.56 | 16/59 |
| Lys | 1.01 ± 0.30 | 10/325 | 0.77 ± 0.25 | 2/322 | 1.14 ± 0.22 | 10/325 | 0.70 ± 0.21 | 1/325 | 0.87 ± 0.28 | 8/325 |
| Tyr | 1.49 ± 1.25 | - | n/a | n/a | 1.88 ± 1.46 | - | 1.24 ± 0.85 | - | 1.79 ± 0.95 | - |

<sup>a</sup>PypKa<sup>12</sup> and ANI-2X<sup>33</sup> predictions were made with the local installed software provided by the authors. DeepKa predictions were obtained from the DeepKa web server.<sup>57</sup> n/a (not available) indicates that the model is unable to make predictions. CER of Tyr is not calculated due to the extremely small test sets (3 Tyr<sup>-</sup>). <sup>b</sup>Our test sets likely overlap with ANI-2X’s training set; removing overlap is impossible as the data in Ref<sup>33</sup> is unpublished.
Ref.<sup>33</sup>

The performance of both the CBtree and GAT correlates with dataset size: lowest RMSE for Glu (largest dataset) and highest RMSE for Cys/Tyr (smallest datasets). However, besides the significant decrease in RMSEs, the CBtree’s CERs for Cys and Lys are reduced by 30% and 65%, respectively, relative to the GAT. This suggests that the CBtree is less sensitive to the dataset size than the GAT.

#### KaML-CBtree and KaML-GAT share common missteps

Since the CBtree and GAT have distinct algorithms, we asked if they make different errors in the p*K*_a_ predictions. To address this question, the model residuals, p*K*_a_(pred)-p*K*_a_(expt), are plotted against each other (Figure 6). For both models, most residuals cluster around zero and spread symmetrically, indicating no systematic errors. However, surprisingly, the residuals of the GAT show a high correlation (PCC of 0.73) with those of the CBtree. This indicates that both models produce errors of similar amplitude and of the same sign for most entries in the dataset. Indeed, ensembling CBtree and GAT predictions failed to reduce RMSEs (data not shown).

**Figure 6:**
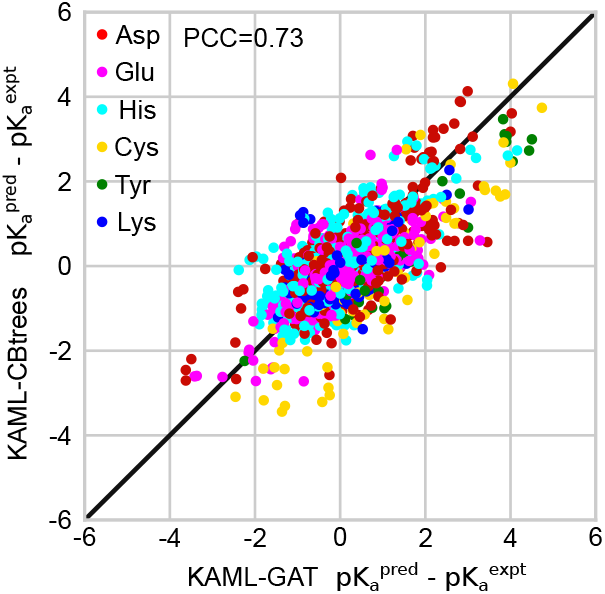
Model residuals are highly correlated between KaML-CBtree and KaML-GAT. For every instance in the test sets the residuals (difference between the predicted and experimental p*K*_a_’s) of the CBtree and GAT are plotted against each other. The solid line is identity.

Close examination of the largest residuals reveals that both models have difficulty predicting the anomalous p*K*_a_’s of mutant SNase. An extreme example is T41D SNase. In the (homology) modeled structure, Asp41 is deeply buried with only hydrophobic sidechains in its surrounding and (accordingly) the predicted p*K*_a_ by either GAT or CBtree is ∼8; however, the experimental value is ∼4. Analogously, the deeply buried Lys132 in A132K SNase has an experimental p*K*_a_ of 10.4, whereas the CBtree and GAT predict 6.9 and 8.9, respectively. This residue is also in a hydrophobic region with only one polar sidechain nearby. Constant pH MD simulations^54^ and experimental evidence^55,56^ suggest that abnormally small p*K*_a_ shifts of deeply buried residues are due to partial unfolding and/or water penetration which allows stabilization of the ionized form.

An example where both models fail to recapitulate large experimental p*K*_a_ shift is Asp118 in N118D SNase (PDB: 5KGU), which has an experimental p*K*_a_ of 7.0, but both GAT and CBtree predict a p*K*_a_ of ∼4.0. Asp118 is in a unique environment: its sidechain is fully buried near positively charged Lys and Arg, while its backbone lies close to the protein surface. We hypothesized that both models overestimate attractive electrostatic contributions stabilizing the charged state; however, the following two examples of large residuals contradict this. Asp75 in barnase (PDB: 1FW7) has an experimental p*K*_a_ of 3.1, while the CBtree and GAT predicts 7.3 and 6.5, respectively. Cys48 in glutathione S-transferase (PDB: 5X79) has an experimental p*K*_a_ of 3.7, while CBtree and GAT predict 7.2 and 8.8, respectively. The microenvironment of Asp75 in barnase and Cys48 in S-transferase share key similarities with Asp118 in N118D SNase: in each case, the titratable sidechain is buried in close proximity to one or two charged sidechains, while the backbone is positioned near the surface. However, the models either under-(Asp118) or overestimate (Asp75 and Cys48) the attractive electrostatics. For edge cases of buried residues in proximity to both solvent and charged sidechains, small conformational changes can significantly alter the local environment of the titratable residue. The effects due to conformational changes are not captured by the ML models.

### Comparison between KaMLs and other models

#### KaML-CBtree significantly outperforms the PB and alternative ML models

The CBtree outperforms the PB method (PyPKa)^12^ and alternative ML models (PKAI, PKAI+,^25^ DeepKa^29,30^), compared to the overall metrics reported (SI Table S9). However, this comparison is not optimal due to the different test sets used. Moreover, the published models^25,29,30,33^ were evaluated using only one test set, making a fair comparison even harder. Furthermore, many models ignore Cys and Tyr, e.g. DeepKa,^29,30^ which results in underestimation of the overall RMSE since Cys and Tyr are associated with larger errors. To ensure a fair comparison among different models, we used our 20 holdout test sets to evaluate the amino acid specific metrics for predicting p*K*_a_’s and protonation states by PypKa,^12^ DeepKa,^30,57^ and ANI-2X.^33^ Note, our test sets likely overlap with the training data of ANI-2X; however, removing overlap is impossible due to unavailability of data in Ref.^33^

KaML-CBtree’s RMSEs and CERs are significantly lower than PypKa and DeepKa for all amino acids (Table 3 and SI Table S10). KaML-CBtree’s RMSEs and CERs are also significantly lower than ANI-2X, except for His, for which ANI-2X’s RMSE is 0.52. Considering the author-reported test RMSE of 0.98 for His^33^ and the significantly higher RMSEs of Asp (1.20) and Glu (0.81) in our tests (Asp and Glu were trained using much larger training datasets), we attribute the unexpectedly low RMSE for His to an overlap between our test sets and ANI-2X’s training set.

Comparison of KaML-GAT’s metrics with those of PypKa and DeepKa shows that KaML-GAT excels at predicting the p*K*_a_’s and protonation states of Asp and Cys, while the performance for Glu is similar to DeepKa which surpasses PypKa (Table 3). Interestingly, KaML-GAT’s RMSE and CER for Lys are lower than PypKa and ANI-2X, which have similar performances; however, DeepKa’s RMSE is 0.1 units and CER is four times lower compared to KaML-GAT. We attribute DeepKa’s excellent performance for Lys to the high accuracy of the GBNeck2-CpHMD titration for Lys^5^ and its significantly larger training data of downshifted p*K*_a_’s compared to KaML-GAT.

Curiously, although KaML-GAT’s RMSE for His is more than 0.24 lower than PypKa and DeepKa, the CER is respectively 3.2 and 2.2 times (Table 3). This performance is related to KaML-GAT’s low Pre and Rec for predicting His^+^ (Figure 5D). The largest residuals for PypKa are from either overestimating extremely low experimental p*K*_a_values or underestimating experimental p*K*_a_values around 6.0. Those instances lead to an increase in RMSE without affecting the CER.

## Concluding Discussion

We developed the shallow decision tree (KaML-CBtree) and graph deep learning (KaML-GAT) models to predict p*K*_a_’s and protonation states of all six titratable amino acids based on a newly curated, significantly expanded experimental p*K*_a_ database PKAD-3 While both KaML-CBtree and KaML-GAT outperform PROPKA3, KaML-CBtree offers more accurate p*K*_a_ and protonation state predictions for all six titratable amino acids. KaML-CBtree’s RMSEs and CERs are also significantly lower than the PB method (PypKa) and ML models trained on the state-of-the-art GBNeck2-CpHMD p*K*_a_’s (DeepKa)^30,57^ and atom-centered quantum potential energies (ANI-2X).^33^

Perhaps the most significant improvement over previous models is the model’s capability of accurately predicting Asp^0^, Glu^0^, Cys^−^, and Lys^0^, which often play important roles in biological functions. In contrast, previous models either fail or are incapable of making predictions. For Asp^0^, PypKa, ANI-2X, and DeepKa produce Rec of 0, 0.09, and 0.12 respectively. For Glu^0^, PypKa, ANI-2X, and DeepKa yield Rec of 0.24, 0, and 0.79, respectively. While ANI-2X and DeepKa are not trained to make predictions for Cys, PypKa gives Rec of 0.16 for Cys^−^. Neither ANI-2X nor PypKa predicts any Lys^0^, i.e. Rec of 0. We suggest that the improved prediction of Asp^0^ and Glu^0^ is not only due to the significant enrichment of highly upshifted p*K*_a_ values, e.g., above 7.5, in the training dataset, but also due to the separate training of acid and base models. The latter allows similar features to make opposite p*K*_a_ contributions, as demonstrated in Figure 4.

Another possible contributor to KaMLs’ improvement over previous models is the expansion of “anomalous” p*K*_a_’s in the training data, many of which are from the Garcia-Moreno lab’s experiments,^40–43^ e.g., Lys100 in N100K SNase, which is deeply buried without nearby electrostatic/h-bond interactions but has a p*K*_a_ downshift of less than 2 units.^45^ To illustrate this point further, a plot of experimental Asp/Glu p*K*_a_’s in PKAD-3 vs. the calculated buried ratios shows that Asp/Glu with highly shifted p*K*_a_’s (e.g., above 7.5) are indeed largely buried; however, many buried Asp/Glu have normal p*K*_a_’s (SI Figure S8). Accurate prediction of these anomalous p*K*_a_’s remains the greatest challenge for structurebased methods such as KaMLs (see earlier discussion of Asp41 in T41D SNase).

One surprising finding is that the performance of the KaML-CBtree well exceeds KaML-GAT, despite having two orders of magnitude smaller parameter space (∼ 8,000 for GAT). Decision trees benefit from simplicity and interpretability, which can lead to more efficient training and better model generalization. Deep learning models are potentially more accurate but require much larger training dataset. The impact of dataset size is evident in KaML-GAT’s performance improvement with the AF2 data augmentation (data not shown), while KaML-CBtree shows no such sensitivity (SI Table S11). Two recent studies^58,59^ found that regularity, i.e., feature distributions that are less skewed and less heavy-tailed, is predictive of neural networks outperforming gradient-boosted tree models for tabular data. Thus, we hypothesized that, in addition to the small dataset, the higher performance of KaML-CBtree may be attributed to the irregularity of features. McElfresh et al.^59^ introduced a feature irregularity parameter as a linear function of specific meta features. Using a large number of datasets and models, they found^59^ that the feature irregularity score ranges from 0 to 7 and tree models outperform neural networks when the irregularity score is greater than 5. When calculating the feature irregularity score using the tree features of the entire dataset, we obtained a value of 6.3, which suggests highly irregular features, thus providing a rationale for why KaML-CBtree outperforms KaML-GAT. On the other hand, the lower performance of KaML-GAT may also be attributed to the limitations of the pretraining data, especially the PHMD549 dataset’s exclusion of Cys and Tyr. This can be seen from the significantly higher RMSEs of Cys and Tyr p*K*_a_’s as compared to those of KaML-CBtree.

The present models have several other limitations. The training and testing datasets for Tyr and Cys are extremely small, leading to larger prediction errors and potentially unreliable model evaluation metrics. Another challenge is related to the class imbalance between the protonated and deprotonated states, which reduces the prediction accuracy for the minority class. For this reason, the recall of His^+^ is significantly lower than that of any other amino acid’s minority protonation state. In our current evaluation of regression models’ classification power, the titrating class is excluded. This leads to an incomplete picture of model performance, particularly for His, as the most probable experimental p*K*_a_’s are near 7. Furthermore, in some applications, the ability to accurately identify titrating residues is crucial for understanding pH-dependent behavior. Despite these limitations, KaML-CBtree demonstrates promising precision and accuracy for predicting protein electrostatics. To enable applications and facilitate further development in the community, we released PKAD-3 and an end-to-end p*K*_a_ prediction tool based on KaML-CBtree.

## Acknowledgment

Financial support by the National Institutes of Health (R35GM148261 and R01CA256557) is acknowledged. We thank Dr. Guy Dayhoff for providing a command line version of the protein disorder prediction tool RIDAO.^60^

## Data Availability

The PKAD-3 database is freely searchable and downloadable at https://database.computchem.org/pkad-3. All training and test sets, the 20 CBtree and GAT models, and the Python program for end-to-end p*K*_a_ predictions using the finalized CBtree model are freely downloadable at https://github.com/JanaShenLab/KaMLs/.

## Supporting Information Available

Supporting Information contains Materials and Methods as well as supplementary tables and figures. Table S1 contains the description of features of the tree models. Table S2 and S3 list regression and classification metrics for the different KaML-trees, respectively. Table S4 gives the performance metrics of the acid/base KaML-CBtree with AF2 augmentation. Table S5 and S6 list amino acid specific regression and classifications metrics for KaML-CBtrees and PROPKA3, respectively. Table S7 gives atom types for KaML-GATs. Table S8 lists regression metrics for the different freezing schemes for KaML-GATs. Table S9 compares the GAT model performance with and without data augmentation or pretraining. Table S10 lists amino acid specific regression and classifications metrics for KaML-GAT. Table S11 gives the author-reported regression metrics for the published p*K*_a_ prediction methods. Table S12 gives regression metrics from the previous ML models evaluated on our test sets. Figure S1 shows the histograms of the p*K*_a_ values in the train/test splits. Figure S2 gives the t-SNE analysis of histidine features at pH 7.5. Figure S3 and S6 show the predicted vss experimental p*K*_a_ values for all 20 test sets for KaML-CBtree and KaML-GAT, respectively. Figure S4 displays the histograms of the p*K*_a_ values of the training dataset with the AF2 augmentation. Figure S5 shows the training and validation loss of KaML-GAT. Figure S6 shows the dependence of ensemble GAT on the number of models in the ensemble. Figure S7 shows the experimental Asp and Glu p*K*_a_’s in PKAD-3 vs. the calculated buried ratios.

## Notes

### Competing Interest Statement

The authors have declared no competing interest.

### Summary of Updates

Fixed a few typos, including a missing dash in the web link and SI figure numbers.

