## Supporting Information for "KaMLs for Predicting Protein p*K*_a_ Values and Ionization States: Are Trees All You Need?"

### Materials and Methods

#### PKAD-3 database

**Constructing PKAD-3 with unique residue identifications.** The current protein p$K_a$ database PKAD-2[S1] contains 1,742 entries associated with 211 proteins. After removing the p$K_a$'s without a specific value, e.g., p$K_a$ >11 or p$K_a$ <2, as well as those of termini, 1,569 entries remain. For these entries, we manually verified the accuracy by reading the source literature. The incorrect p$K_a$ values and residue IDs were corrected. For example, the p$K_a$ of Glu172 in the WT xylanase (associated with PDB entry 1XNB) was corrected from 7.2 to 6.7. The residues that are not resolved in the recorded PDB file (e.g., Glu142 on chain A of the PDB file 1STN) were removed. The cleaned PKAD-2 contains 1,286 entries, which are associated with 763 p$K_a$'s of 615 residues in 113 proteins.

Next, we conducted an exhaustive search for additional experimental p$K_a$'s in peer-reviewed articles published before April 2023. First, we applied keyword p$K_a$ in the title and/or abstract to identify reference articles on the PubMed website. Next, we manually searched for p$K_a$ values in the articles. If the authors of the article associated the p$K_a$'s with an X-ray crystal structure, we recorded its PDB ID. If such information was unavailable, we utilized the UniProt ID of the protein to identify an X-ray structure from the RCSB PDB website.[S2] Additionally, we included 79 p$K_a$'s of mutated Asp, Glu, and Lys of the WT or stabilized constructs (PHS and $\Delta$+PHS) of staphylococcal nuclease (SNase). These p$K_a$'s (documented in Ref.[S3]) were derived from thermodynamic stability measurements conducted by the Garcia-Moreno lab and released in the 2009 blinded protein p$K_a$ prediction challenge.[S4] The uncertainty of these p$K_a$'s was estimated to be 0.2–0.5 units,[S5] similar to the p$K_a$'s determined by NMR shifts.[S6] We designated the cleaned and significantly expanded database as PKAD-3 (Table 1) in maintaining continuity with the original database name PKAD[S7] and PKAD-2.[S1]

To make PKAD-3 ML ready and prevent data leakage (across training, validation, and test datasets) caused by the different numbering of residues in different PDB files, entries are annotated by the UniProt ID of the protein and PDBrenum[S8] was used to renumber the residue IDs according to the UniProt sequence. Thus, each residue is identified as UniProt ID+Uni_resid. Note that previous ML model developments[S9–S12] have a data leakage issue, in which some residues with different IDs in different PDB files were treated as distinct residues, while in fact they are the same. Below we discuss other steps undertaken to ensure the data is accurate and ML ready.

**Mutant proteins and multiple conformational states.** Another potential source of data leakage in training ML models is related to annotation of mutant entries. PKAD-3 contains 186 p$K_a$ entries associated with 97 residues in mutants of 6 proteins (Uniprot IDs: P0AEG4, P03069, P00720, P00644, P00282, P09850). Additionally, 100 p$K_a$ entries are associated with 40 residues in 8 proteins (Uniprot IDs: P10599, P68871, P69905, P02945, P00392, Q72F37, P00760, P80028) that have multiple conformational states. These entries were verified to ensure that the X-ray structures are correctly associated with the protein or the conformational state of interest.

To prevent leakage across the training, validation, test datasets, mutant proteins are annotated as 'Uniprot ID-mutation or conformational state', e.g., P09850-Q7E for mutant Q7E; P10599-1 and P10599-2 for two conformational states. A special case is SNase, which has three background constructs (WT, PHS, and $\Delta$+PHS) for engineered mutations. In this case, we use, for example, P00644-V66D, P00644-WT-V66D, and P00644-PHS-V66D to denote the $\Delta$+PHS, WT, and PHS constructs, respectively.

**Entries associated with the same residue but different p$K_a$'s.** We verified the p$K_a$ entries associated with the same protein residue but differ by more than 1 unit. In certain instances, the large p$K_a$ difference is due to a mutation in the protein, in which case, we verified that the mutant is represented by an appropriate X-ray structure or a computational model (see Modeling of mutant structures). In other instances, the large p$K_a$

difference is due to a conformational change, in which case we verified that the conformational state discussed in the literature is indeed represented by the recorded PDB entry. For example, in the oxidized state of thioredoxin (PDB entry 1ERU), the p$K_a$ of Asp26 is 8.1, while in the reduced state the p$K_a$ is 9.9 (PDB entry 1ERT).[S13] In some instances, the large p$K_a$ difference cannot be "reconciled". For example, Glu172 has two measured p$K_a$'s, 7.2 (PDB entry 1XNB) and 2.8 (PDB entry 3VZO). In the latter case, a nearby residue Glu78 is covalently linked to the substrate and another nearby residue is mutated, which likely results in the large p$K_a$ downshift of Glu172. Since ligand binding is not accounted for by the ML models, the p$K_a$ entry 2.8 was removed. For Asp85 in bacteriorhodopsin, the reported p$K_a$'s are 2.6 (PDB: 1UCQ) and 7.2 (PDB: 1IW6). We removed the former entry, since retinol in the 1UCQ structure forms a salt bridge with Asp85. For Asp102 in trypsin, the reported p$K_a$'s are 1.4 (PDB: 2TGA)[S14] and 6.8 (PDB: 1C5Q).[S15] We removed latter p$K_a$, as it was later assigned to the coupling partner His57.[S14]

**Excluded p$K_a$ values.** The p$K_a$'s of residues interacting with HEME or ligand (e.g., PDB: 3VZO, Glu172) were excluded, as the ML models do not account for HEME or ligand. Furthermore, we removed Glu325 of LacY (PDB: 2V8N) with a reported p$K_a$ of 10.5.[S16] It is the only Glu with such an extreme (6.3 unit) p$K_a$ shift relative to the model, and the value itself is suspicious. Sugar transport of LacY is coupled to proton co-transport involving Glu325. However, a p$K_a$ of 10.5 makes de-protonation unlikely. Furthermore, the p$K_a$ of Glu325 has been suggested to depend on the conformation of LacY.[S16] In light of these considerations, we decided to exclude Glu325 of LacY in PKAD-3.

#### Preparation of structures for ML training

Protein structure files were downloaded from the PDB. The PDBfixer function from OpenMM[S17] was used to add missing atoms/residues and remove ligands and waters. The PDBrenum software[S8] was used to renumber the residue IDs according to the UniProt sequence[S18] to ensure that each residue together with the protein Uniprot ID is assigned a unique identification. For multimer proteins, the monomer that contains the residue with experiment p$K_a$ was retained. Unless X-ray structure is available, the structure of a mutant protein was modeled by SWISS-MODEL[S19] using the X-ray structure of the WT (or the appropriate background construct in the case of SNase) as the template.

**Data augmentation using AlphaFold2 models.** AlphaFold2 (AF2)[S20] predictions were made using the colabfold_batch program from the ColabFold v1.5.5[S21] docker image. The multiple sequence alignment (MSA) sampling was carried out on the MMSeqs2[S22] server and the results were downloaded for the remainder of the calculations. MSA sampling used 32 cluster centers and 64 extra sequences. We used 32 random seeds and enabled dropout at the default rate for more varied structures. Lastly, we recycled each structure three times, since higher recycling did not increase the template-modeling (pTM) scores. The default parameters were used for all other ColabFold parameters.

#### Development of KaML-trees and KaML-GATs

**Data splitting protocol.** The "StratifiedGroupKFold" strategy adapted from sklearn[S23] is applied to randomly sample 10% of PKAD-3 as the hold-out test set. Employing the same strategy, the remaining 90% is divided into training and validation sets in a 9:1 ratio. To ensure that the models are evaluated with statistical uncertainties, the data splitting process will be repeated 20 times to generate 20 independent ML models. The same training and test data are used for KaML-trees and KaML-GATs.

A unique group is defined as a combination of UniProt ID+Uni_resid. Stratification was performed on the p$K_a$'s to ensure a similar distribution between the training and test datasets. Specifically, for each amino acid, we manually binned the unique residues based on their p$K_a$'s. The bin width was defined such that residues with different p$K_a$'s can be separated while most of the bins can be further split into training, validation, and test sets (SI Figure S1).

**Feature engineering for the KaML-trees.** To describe the local environment of the titratable residue, we adapted six types of features used by the tree models for cysteine ligandability predictions:[S24]

- basic characteristics of the titratable residue;
- solvent accessibility;
- potential hydrogen bond (h-bond) interactions;
- potential salt-bridge interactions;
- propensity of local disorder;
- secondary structure;

A complete list of the features and explanations is given in Table S1.

**Training the KaML-trees.** For training KaML-trees, the labels were the absolute p$K_a$'s, and the training loss function was the mean squared error (MSE). Model training was repeated 20 times with 20 sets of training/validation and test datasets split as discussed above. 10-fold cross-validation was employed. The open-source Python libraries scikit-learn,[S23] Pycaret,[S25] and CatBoost[S26] were employed for model training and hyperparameter optimization. The latter utilized the random search algorithm with 5000 iterations. The optimized hyperparameters were used to train a final model based on the entire PKAD-3, and the model was saved for user deployment.

**AF2 structure augmentation.** Out of the 32 AF2 predicted structures for each protein, we only added those in a similar conformation as the X-ray structure. The conformational similarity was defined by the numerical features of the tree models. First, all numerical features were min-max normalized. Next, a similarity score S(f1,f2) was calculated as the maximum of the squared differences.

$$S(f_1, f_2) = \max_{1 \leq i \leq N_{\times features}} \{(f_{1i} - f_{2i})^2, \} \tag{1}$$

where f1 and f2 refer to the features of the X-ray and AF2 structures, respectively. We tested different augmentation schemes defined by the threshold of p$K_a$ shifts ($> 1$ or $> 2$), the similarity cut-off ($0.04$ or $0.01$) as well as a maximum number of AF2 structures to add (at most $10$ or all within the similarity cut-off).

**Calculation of data irregularity.** To investigate feature irregularity, we followed the approach used by McElfresh and coworkers.[S27] The Python package PyMFE[S28] was used to extract meta-features from the tree features of the entire PKAD-3 dataset. From the meta-features, an irregularity score (6.3) was calculated as weighted sum of the smallest eigenvalue of the covariance matrix (weight of -0.33), the skewness of the standard deviation (weight of 0.23), the skewness of the range (weight of 0.22), the interquartile range of the harmonic mean (weight 0f 0.21), and the standard deviation of the kurtosis (weight of 0.21).

**Graph representation of the protein structures.** The Python package Graphein[S29] was used to build graphs that represent the titratable residues. For computational efficiency, similar to the 20-Å cubic box used in DeepKa,[S30,S31] the protein structure was truncated using BioPandas[S32] as a sphere of 10-Å radius around the most relevant atoms in the titratable sidechain: OD1/OD2 for Asp, OE1/OE2 for Glu, ND1/NE2 for His, SG for Cys, NZ for Lys, and OH for Tyr. Within this sphere, all atoms were considered nodes. The graph incorporates 7 edge types based on pairwise atomic distances. Bonded edges connect atom pairs within 2.2 Å, while non-bonded edges link atoms separated by 2.2-3.5 Å. A bonded edge has 3 subtypes: single, double (based on the distance cutoffs[S33]), and ring (edge within a cycle). A non-bonded edge (with a cutoff of 3.5 Å) has 4 subtypes:

hydrophobic (between Ala, Val, Leu, Ile, Met, Phe, Trp, Pro, and Tyr), aromatic (between Phe, Trp, His, and Tyr), ionic (between Arg, Lys, His, Asp, and Glu), and hydrogen bond (h-bond). A h-bond is searched between sidechain h-bond donors: ARG (NE, NH1, NH2), ASN (ND2), GLN (NE2), HIS (ND1, NE2), LYS (NZ), SER (OG), THR (OG1), TRP (NE1), or TYR (OH) and h-bond acceptors: ASN (OD1), ASP (OD1, OD2), GLN (OE1), GLU (OE1, OE2), or HIS (ND1, NE2). Currently, the backbone atoms and Cys sulfur are not considered in building h-bond edges by Graphein,[S29] which is a limitation and will be addressed in our future work.

We devised a 42-digit-vector node embedding scheme similar to the one used in the CNN model for predicting cysteine ligandabilities.[S24]

- 24-digit one-hot encoding for the atom type of the CHARMM c22 force field[S34] (see SI Table S6);
- 1 integer to indicate if the atom belongs to the titratable residue (0: no; 1: yes);
- 7-digit one-hot for amino acid: Asp, Glu, Cys, Tyr, His, Lys, or other;
- 1 integer for the atom hybridization state;
- 1 integer for the number of bonded heavy atoms;
- 1 integer for the number of hetero atoms bonded;
- 1 float for the atom partial charge from the CHARMM c22 force field;[S34]
- 1 float for the model p$K_a$ (Asp 3.7, Glu 4.2, His 6.5, Cys 8.5, Tyr 10.1, and Lys 10.4);
- 5-digit one-hot to indicate hydrophobic, aromatic, h-bond acceptor, h-bond donor, or ring.

**Architecture and hyperparameters of the KaML-GATs.** The PyTorch Geometric (PyG) library[S35] was used to build the KaML-GATs (Figure 5 in the main text). The KaML-GAT contains 3 one-head convolutional (conv) layers, each having 42 channels based on the graph attention operator.[S36] The attention coefficient between the $i^{th}$ and the $j^{th}$ node can be expressed as

$$\alpha_{ij} = \frac{\exp(LeakyReLU(\overrightarrow{a}^T[W\overrightarrow{h}_i||W\overrightarrow{h}_j]))}{\sum_{k\in N_i}\exp(LeakyReLU(\overrightarrow{a}^T[W\overrightarrow{h}_i||W\overrightarrow{h}_k]))}, \tag{2}$$

where $\overrightarrow{a}^T$ is the attention coefficient, W is the linear transformation weight, and || denotes the aggregation function. The LeakyReLU is an activation function defined as:

$$LeakyRelu(x) = max(0, x) + 0.2 * min(0, x). \tag{3}$$

A hyperbolic tangent (tanh) layer is added after each conv layer to add non-linearity to the model. A global mean pooling layer is applied to the output of the last conv layer to represent the whole graph embedding. The pooling results are flattened to a vector of length 42 and fed to the fully connected (FC) layer. The FC layer is made up of 2 hidden layers containing 32 and 16 neurons. The rectified linear unit (ReLU) activation function functions is applied to all the hidden layers. Finally, 1 FC layer containing 1 neuron with a linear activation function is used to make prediction. Batch normalization is applied to all the layers except the last FC layer. A training batch size of 64 and a learning rate of 0.0005 was used. A dropout rate of 0.2 was applied to both conv and FC layers. The Adam optimizer was used for minimizing training loss MSE. To prevent over-fitting, early stopping was applied if the validation loss stops decreasing in the next 50 epochs with a tolerance of 0.1. The total number of parameters in KaML-GAT is $\sim 8000$.

**Training the KaML-GATs.** The Pytorch API[S37] was used to implement model training. Training labels were the experimental (in training) or calculated (in pre-training) p$K_a$ shifts relative to the model values, while in analysis, the shifts were converted back to the p$K_a$'s. Model pre-training utilized PHMD549,[S31] a dataset comprised of Asp, Glu, His, and Lys p$K_a$'s calculated by the pH-replica exchange[S38] GBNeck2-CpHMD[S39] simulations. After removing proteins in the PKAD-3 database, there were 26,252 p$K_a$'s of 25,912 residues in 535 proteins. Note, this dataset does not contain p$K_a$'s of Cys and Tyr, and the highly shifted p$K_a$'s may not be accurate due to the short simulation time.[S31] During pre-training, the model weights were initialized using the Glorot algorithm.[S40]

Once pre-training was completed, different layer freezing schemes were tested (see main text) before training on the experimental p$K_a$ shifts. For each data split, 10 GATs were trained using different 9:1 train-CV splits independent of each other. The models with the validation RMSE $<$ 1.2 were selected to form an ensemble GAT for hold-out testing.

#### A web portal for the PKAD-3 database

We implemented a web portal (https://database.computchem.org/pkad-3). Each entry contains the PDB ID, chain ID, residue type, residue ID (in the PDB file), p$K_a$ value, Uniprot_id, Uniprot_resid. The Note column contains information about mutation and/or protein conformational state. Hyperlinks for the cited references are also included. Users can search the PKAD-3 database by specific columns (e.g., PDB, Chain) and apply criterion to filter p$K_a$ values that meet certain criteria (e.g., $> 7$). Users can also select specific rows from their search results. The user selection, the entire search result, or the complete database can be downloaded as a csv file.

### Supplemental Tables

Table S1: Features used for the decision tree ML models

| Feature name | Description |
|---|---|
| | **Characteristics of the titratable residue** |
| res_type | residue type |
| model_p$K_a$ | p$K_a$ of model compound in solution |
| charge | residue charge at physiological pH |
| n_sc_C | number of side chain carbon atom |
| | **Solvent Accessibility** |
| buried_ratio | 1 - SASA(X)/SASA(model) |
| d0/1_polar | distance from the side chain COM to the first/second nearest polar atom |
| d0/1_nonpolar | distance from the side chain COM to the first/second nearest nonpolar atom |
| n_polar_5/10/15 | number of polar atoms within 5/10/15Å of the side chain COM |
| n_nonpolar_5/10/15 | number of nonpolar atoms within 5/10/15Å of the side chain COM |
| n_hv_6/9/12/15 | number of heavy atoms within 6/9/12/15Å of the side chain COM |
| | **Potential h-bond interactions** |
| d0/1_hbond_O | distance from the side chain COM to the first/second nearest Ser/Thr/Tyr side chain O |
| d0/1_hbond_N | distance from the side chain COM to the first/second nearest Asn/Gln/Trp side chain N |
| d0/1_hbond_H | distance from the side chain COM to the first/second nearest His side chain N |
| d0/1_C_SG | distance from the side chain COM to the first and second nearest Cys side chain SG |
| d0/1_bb_NH | distance from the side chain COM to the first/second nearest backbone N |
| | **Potential electrostatic interactions** |
| d0/1_neg_O | distance from the side chain COM to the first/second nearest Asp/Glu side chain O |
| d0/1_pos_N | distance from the side chain COM to the first/second nearest Arg/Lys side chain N |
| | **Protein flexibility** |
| flexibility | predicted by RIDAO[S41] on a 30-amino acid stretch centered at the titratable residue. |
| | **Protein secondary structure** |
| rss | secondary structure of the ROI |
| rssp2/4 | secondary structure of the residue at +2/+4 position |
| rssm2/4 | secondary structure of the residue at -2/-4 position |
| | **Potential backbone h-bond interactions** |
| DA1/2 | Distance from the backbone N of the titratable residue to the nearest/second nearest backbone O of any other residue |
| DD1/2 | Distance from the backbone O of the titratable residue to the nearest/second nearest backbone N of any other residue |

[a] SASA (solvent accessible surface area) is calculated by DSSP.[S42] Model refers to the penta-peptide AAXAA, where X = ROI. A command line version of RIDAO[S41] was used.

Table S2: Overall performance comparison of the different KaML-trees for predicting p$K_a$values

| | CB | | | ET | | | GB | RF | XGB |
|---|---|---|---|---|---|---|---|---|---|
| | all | acid | base | all | acid | base | all | all | all |
| PCC | $0.94 \pm 0.02$ | $0.88 \pm 0.03$ | $0.93 \pm 0.03$ | $0.94 \pm 0.02$ | $0.86 \pm 0.03$ | $0.93 \pm 0.03$ | $0.94 \pm 0.02$ | $0.94 \pm 0.02$ | $0.93 \pm 0.02$ |
| RMSE | $0.77 \pm 0.09$ | $0.77 \pm 0.11$ | $0.76 \pm 0.09$ | $0.80 \pm 0.10$ | $0.82 \pm 0.13$ | $0.74 \pm 0.09$ | $0.81 \pm 0.07$ | $0.81 \pm 0.08$ | $0.86 \pm 0.09$ |
| MAXE | $3.47 \pm 0.39$ | $3.25 \pm 0.53$ | $2.60 \pm 0.70$ | $3.70 \pm 0.78$ | $3.61 \pm 1.01$ | $2.46 \pm 0.74$ | $3.79 \pm 0.49$ | $3.66 \pm 0.58$ | $4.08 \pm 0.85$ |

[a]The metrics of the CBtree are shown in bold font. All models were trained on the non-augmented dataset.

Table S3: Overall performance comparison of the different KaML-trees for predicting protonation states at pH 7[a]

| | CB | | | ET | | | GB | RF | XGB |
|---|---|---|---|---|---|---|---|---|---|
| | all | acid | base | all | acid | base | all | all | all |
| $\text{Rec}^{\text{prot}}$ | **0.93** | 0.82 | 0.96 | 0.90 | 0.88 | 0.91 | 0.93 | 0.93 | 0.93 |
| $\text{Pre}^{\text{prot}}$ | **0.97** | 0.91 | 0.99 | 0.95 | 0.88 | 0.98 | 0.96 | 0.96 | 0.91 |
| $\text{Rec}^{\text{dep}}$ | **0.98** | 0.99 | 0.99 | 0.94 | 0.99 | 0.97 | 0.99 | 0.99 | 0.98 |
| $\text{Pre}^{\text{dep}}$ | **0.99** | 0.99 | 0.95 | 0.98 | 0.99 | 0.88 | 0.98 | 0.98 | 0.98 |
| CER | 46/2635 | 34/2099 | 12/536 | 64/2638 | 30/2061 | 34/577 | 46/2578 | 48/2533 | 72/2606 |

[a]The metrics of the CBtree are shown in bold font. All models were trained on the non-augmented dataset.

Table S4: Performance metrics of the acid/base KaML-CBtree with AF2 structure augmentation

| | w/o aug | 2.0,0.04,all | 2.0,0.04,10 | 2.0,0.01,all | 2.0,0.01,10 | 1.0,0.04,all | 1.0,0.04,10 | 1.0,0.01,all | 1.0,0.01,10 |
|---|---|---|---|---|---|---|---|---|---|
| PCC | $0.94 \pm 0.02$ | $0.94 \pm 0.02$ | $0.94 \pm 0.02$ | $0.94 \pm 0.02$ | $0.94 \pm 0.02$ | $0.93 \pm 0.02$ | $0.94 \pm 0.02$ | $0.94 \pm 0.02$ | $0.94 \pm 0.02$ |
| RMSE | $0.77 \pm 0.09$ | $0.84 \pm 0.10$ | $0.81 \pm 0.09$ | $0.82 \pm 0.08$ | $0.83 \pm 0.10$ | $0.85 \pm 0.12$ | 0.840.11 | $0.81 \pm 0.08$ | $0.83 \pm 0.10$ |
| MAXE | $4.1 \pm 0.85$ | $3.60 \pm 0.55$ | $3.8 \pm 0.38$ | $3.61 \pm 0.42$ | $3.8 \pm 0.5$ | $3.7 \pm 0.5$ | $3.8 \pm 0.4$ | $3.8 \pm 0.2$ | $3.7 \pm 0.4$ |
| $\text{Pre}^{\text{prot}}$ | 0.91 | 0.90 | 0.90 | 0.91 | 0.92 | 0.88 | 0.88 | 0.91 | 0.91 |
| $\text{Pre}^{\text{dep}}$ | 0.94 | 0.94 | 0.94 | 0.93 | 0.93 | 0.94 | 0.94 | 0.95 | 0.94 |
| $\text{Rec}^{\text{prot}}$ | 0.74 | 0.75 | 0.76 | 0.74 | 0.74 | 0.77 | 0.78 | 0.75 | 0.75 |
| $\text{Rec}^{\text{dep}}$ | 0.94 | 0.93 | 0.93 | 0.93 | 0.93 | 0.93 | 0.93 | 0.94 | 0.94 |
| CER | 46/2635 | 51/2635 | 46/2635 | 41/2635 | 55/2635 | 59/2635 | 57/2635 | 48/2635 | 53/2635 |

Models were trained with augmented dataset, column labels indicate augmentation parameters ($\Delta$p$K_a$, similarity, number of structures).

Table S5: Performance metrics of KaML-CBtree for individual residue types

| | Asp | Glu | His | Cys | Lys | Tyr |
|---|---|---|---|---|---|---|
| PCC | $0.86 \pm 0.06$ | $0.84 \pm 0.04$ | $0.51 \pm 0.19$ | $0.61 \pm 0.52$ | $0.80 \pm 0.16$ | $0.39 \pm 0.85$ |
| RMSE | $0.75 \pm 0.17$ | $0.60 \pm 0.07$ | $0.85 \pm 0.14$ | $1.50 \pm 0.60$ | $0.70 \pm 0.21$ | $1.24 \pm 0.85$ |
| Protonation state prediction at pH 7.0 | | | | | | |
| $\text{Prec}^{\text{prot}}$ | 0.95 | 1.0 | 1.0 | 0.68 | 0.99 | – |
| $\text{Rec}^{\text{prot}}$ | 0.77 | 0.81 | 0.37 | 0.65 | 1.0 | – |
| $\text{Prec}^{\text{dep}}$ | 0.99 | 0.99 | 0.95 | 0.73 | 1.0 | – |
| $\text{Rec}^{\text{dep}}$ | 0.99 | 1.0 | 1.0 | 0.76 | 0.80 | – |
| CER | 13/916 | 5/1076 | 11/209 | 13/68 | 1/325 | |
| Protonation state prediction at pH 7.5 | | | | | | |
| $\text{Prec}^{\text{prot}}$ | 0.91 | 1.0 | 1.0 | 0.75 | 0.99 | – |
| $\text{Rec}^{\text{prot}}$ | 0.67 | 0.56 | 0.18 | 0.38 | 1.0 | – |
| $\text{Prec}^{\text{dep}}$ | 0.99 | 0.99 | 0.98 | 0.71 | 1.0 | – |
| $\text{Rec}^{\text{dep}}$ | 0.99 | 1.0 | 1.0 | 0.93 | 0.80 | – |
| CER | 6/896 | 7/1068 | 9/390 | 12/43 | 2/315 | |
| Number of protonated/deprotonated states in the test set data[a] | | | | | | |
| $N^{\text{prot}}$ (pH 7.0) | 62 | 40 | 91 | 31 | 315 | 34 |
| $N^{\text{dep}}$ (pH 7.0) | 888 | 1032 | 282 | 36 | 10 | 3 |
| $N^{\text{prot}}$ (pH 7.5) | 24 | 22 | 25 | 20 | 313 | 34 |
| $N^{\text{dep}}$ (pH 7.5) | 892 | 1054 | 417 | 40 | 12 | 3 |

[a] The number of protonated/deprotonated states at pH 7 or 7.5 calculated from the true (experimental) p$K_a$ values. Protonation state is defined by the protonation probability: protonated if Prob $> 0.75$, i.e., p$K_a < 6.52$ at pH 7 or p$K_a < 7$ at pH 7.5; deprotonated if Prob $< 0.25$, i.e., p$K_a > 7.02$ at pH 7 or p$K_a > 7.98$ at pH 7.5.

Table S6: Performance metrics of PROPKA for individual residue types

| | Asp | Glu | His | Cys | Lys | Tyr |
|---|---|---|---|---|---|---|
| PCC | $0.64 \pm 0.16$ | $0.69 \pm 0.12$ | $0.45 \pm 0.23$ | $0.12 \pm 0.75$ | $0.75 \pm 0.16$ | $-$ |
| RMSE | $1.12 \pm 0.16$ | $1.02 \pm 0.23$ | $1.03 \pm 0.25$ | $3.58 \pm 0.81$ | $0.80 \pm 0.22$ | $1.67 \pm 1.80$ |
| Classification at pH 7.0[a] | | | | | | |
| $\text{Pre}^{\text{prot}}$ | $0.84$ | $0.67$ | $0.6$ | $0.47$ | $0.98$ | $-$ |
| $\text{Rec}^{\text{prot}}$ | $0.28$ | $0.93$ | $0.12$ | $1.0$ | $0.99$ | $-$ |
| $\text{Pre}^{\text{dep}}$ | $0.97$ | $0.99$ | $0.85$ | $NaN$ | $0.0$ | $-$ |
| $\text{Rec}^{\text{dep}}$ | $0.99$ | $0.98$ | $0.98$ | $0.0$ | $0.0$ | $-$ |
| CER | $31/907$ | $21/1045$ | $47/303$ | $35/66$ | $6/315$ | |
| Classification at pH 7.5[b] | | | | | | |
| $\text{Pre}^{\text{prot}}$ | $0.56$ | $0.66$ | $0.0$ | $0.44$ | $0.98$ | $-$ |
| $\text{Rec}^{\text{prot}}$ | $0.28$ | $1.0$ | $0.0$ | $1.0$ | $0.99$ | $-$ |
| $\text{Pre}^{\text{dep}}$ | $0.99$ | $1.0$ | $1.0$ | $NaN$ | $0.50$ | $-$ |
| $\text{Rec}^{\text{dep}}$ | $0.99$ | $0.99$ | $1.0$ | $0.0$ | $0.29$ | $-$ |
| CER | $17/908$ | $11/1041$ | $29/434$ | $36/64$ | $7/315$ | - |

[a]Classification of protonation states uses the protonation probability given a predicted p$K_a$, 0–0.25 (or p$K_a > 7.5$ ) for deprotonated and 0.75–1 (or p$K_a < 6.5$) for protonated. Residues with experimental p$K_a$ outside of this ranges were ignored. [b] Classification of protonation states uses the protonation probability given a predicted p$K_a$, 0–0.25 (or p$K_a > 8$ ) for deprotonated and 0.75–1 (or p$K_a$<7) for protonated, Critical error rate (CER) refers to the percentage cases of miss-classifying 'protonated' as 'deprotonated' or vice versa.

Table S7: CHARMM22 atom types

| Atom type | Description |
|---|---|
| C | Carbonyl C, peptide backbone |
| CA | Aromatic C |
| CT1 | Aliphatic sp3 C for CH |
| CT2 | Aliphatic sp3 C for $CH_2$ |
| CT3 | Aliphatic sp3 C for $CH_3$ |
| CPH1 | His CG and CD2 carbons |
| CPH2 | His CE1 carbon |
| CPT | Trp C between rings |
| CY | Trp C in pyrrole ring |
| CP1 | Tetrahedral C (Pro CA) |
| CP2 | Tetrahedral C (Pro CB/CG) |
| CP3 | Tetrahedral C (Pro CD) |
| CC | Carbonyl C, Asn, Asp, Gln, Glu, cter, ct2 |
| N | Pro N |
| NR3 | Charged his ring N |
| NH1 | Peptide N |
| NH2 | Amide N |
| NH3 | Ammonium N |
| NC2 | Guanidinium N |
| NY | Trp N in pyrrole ring |
| O | Carbonyl O |
| OC | Carboxylate O |
| OH1 | Hydroxyl O |
| S | Sulphur |

Table S8: Performance metrics of the pre-trained ensemble KaML-GAT model with different layer freezing schemes[a]

| | freeze0 | freeze1 | freeze2 |
|---|---|---|---|
| PCC | $0.93 \pm 0.02$ | $0.92 \pm 0.03$ | $0.92 \pm 0.04$ |
| RMSE | $0.90 \pm 0.08$ | $0.92 \pm 0.09$ | $0.96 \pm 0.12$ |
| MAXE | $3.74 \pm 0.49$ | $3.74 \pm 0.54$ | $3.90 \pm 0.76$ |
| $\text{Rec}^{\text{prot}}$ | $0.92 \pm 0.06$ | $0.93 \pm 0.06$ | $0.89 \pm 0.07$ |
| $\text{Pre}^{\text{prot}}$ | $0.92 \pm 0.05$ | $0.91 \pm 0.07$ | $0.90 \pm 0.07$ |
| $\text{Rec}^{\text{dep}}$ | $0.98 \pm 0.01$ | $0.98 \pm 0.01$ | $0.98 \pm 0.01$ |
| $\text{Pre}^{\text{dep}}$ | $0.98 \pm 0.01$ | $0.98 \pm 0.01$ | $0.98 \pm 0.02$ |
| CER | $70/2822$ | $75/2822$ | $94/2822$ |

[a]Freeze0, 1, 2 indicates freezing none, the first one, and the first two GAT layers after pretraining.

Table S9: Effect of data augmentation and pretraining on the performance of the GAT model

| Method | RMSE | PCC |
|---|---|---|
| PT + DA | 0.84 | 0.95 |
| DA only | 0.95 | 0.94 |
| PT only | 0.93 | 0.93 |
| No PT/DA | 0.90 | 0.94 |

Model comparison is based on one training/test split.

Table S10: Performance metrics of the ensemble KaML-GAT for different residue types

| | Asp | Glu | His | Cys | Lys | Tyr |
|---|---|---|---|---|---|---|
| PCC | $0.82 \pm 0.08$ | $0.73 \pm 0.07$ | $0.45 \pm 0.18$ | $0.59 \pm 0.56$ | $0.68 \pm 0.21$ | – |
| RMSE | $0.88 \pm 0.17$ | $0.76 \pm 0.08$ | $0.86 \pm 0.18$ | $1.81 \pm 0.56$ | $0.87 \pm 0.28$ | $1.79 \pm 0.95$ |
| $Pre^{prot}$ | 0.95 | 1.00 | 0.25 | 0.58 | 0.97 | – |
| $Rec^{prot}$ | 0.75 | 0.52 | 0.13 | 0.97 | 1.00 | – |
| $Pre^{dep}$ | 1.00 | 0.99 | 0.92 | 0.85 | 0.00 | – |
| $Rec^{dep}$ | 1.00 | 1.00 | 0.94 | 0.47 | 0.00 | – |

Table S11: Overall performance of the PB and ML models

| Method | RMSE |
|---|---|
| PypKa[S43] | $0.82^{a}$ |
| PKAI[S12] | $1.15^{a}$ |
| PKAI+[S12] | $0.98^{a}$ |
| DeepKa[S30,S31] | $1.05^{a,b}$ |
| ANI-2X[S10] | n/a$^{c}$ |
| KaML-CBtree | 0.77±0.07 |
| KaML-GAT | 0.90±0.08 |

$^{a}$ Metrics of alternative models are taken from the references. $^{b}$ CYS and TYR are not included in the RMSE. $^{c}$ The overall RMSE is not given in Ref.[S10]

Table S12: Classification metrics of PypKa, DeepKa, and ANI-2X for individual residue types

| | Asp | Glu | His | Cys | Lys | Tyr |
|---|---|---|---|---|---|---|
| | | | PypKa | | | |
| MAXE | $6.24 \pm 1.3$ | $2.81 \pm 0.74$ | $3.14 \pm 1.49$ | $4.34 \pm 1.68$ | $2.78 \pm 0.72$ | $1.92 \pm 1.59$ |
| $Pre^{prot}$ | $NaN$ | 1.0 | 0.94 | 0.50 | 0.96 | – |
| $Rec^{prot}$ | 0.00 | 0.24 | 0.72 | 1.00 | 1.00 | – |
| $Pre^{dep}$ | 0.94 | 0.98 | 0.97 | 1.00 | $NaN$ | – |
| $Rec^{dep}$ | 1.00 | 1.00 | 0.99 | 0.16 | 0.00 | – |
| | | | ANI-2X | | | |
| MAXE | $4.44 \pm 0.23$ | $4.05 \pm 0.47$ | $1.85 \pm 0.66$ | – | $3.86 \pm 0.55$ | $2.52 \pm 1.71$ |
| $Pre^{prot}$ | 1.00 | $Nan$ | 1.00 | – | 0.96 | – |
| $Rec^{prot}$ | 0.09 | 0.0 | 0.93 | – | 1.00 | – |
| $Pre^{dep}$ | 0.94 | 0.96 | 0.98 | – | $NaN$ | – |
| $Rec^{dep}$ | 1.00 | 1.00 | 1.00 | – | 0.00 | – |
| | | | DeepKa | | | |
| $Pre^{prot}$ | 0.86 | 1.00 | 0.75 | – | 1.00 | – |
| $Rec^{prot}$ | 0.12 | 0.79 | 0.65 | – | 1.00 | – |
| $Pre^{dep}$ | 0.95 | 0.99 | 0.96 | – | 0.86 | – |
| $Rec^{dep}$ | 1.00 | 1.00 | 0.98 | – | 0.86 | – |

#### Supplemental Figures

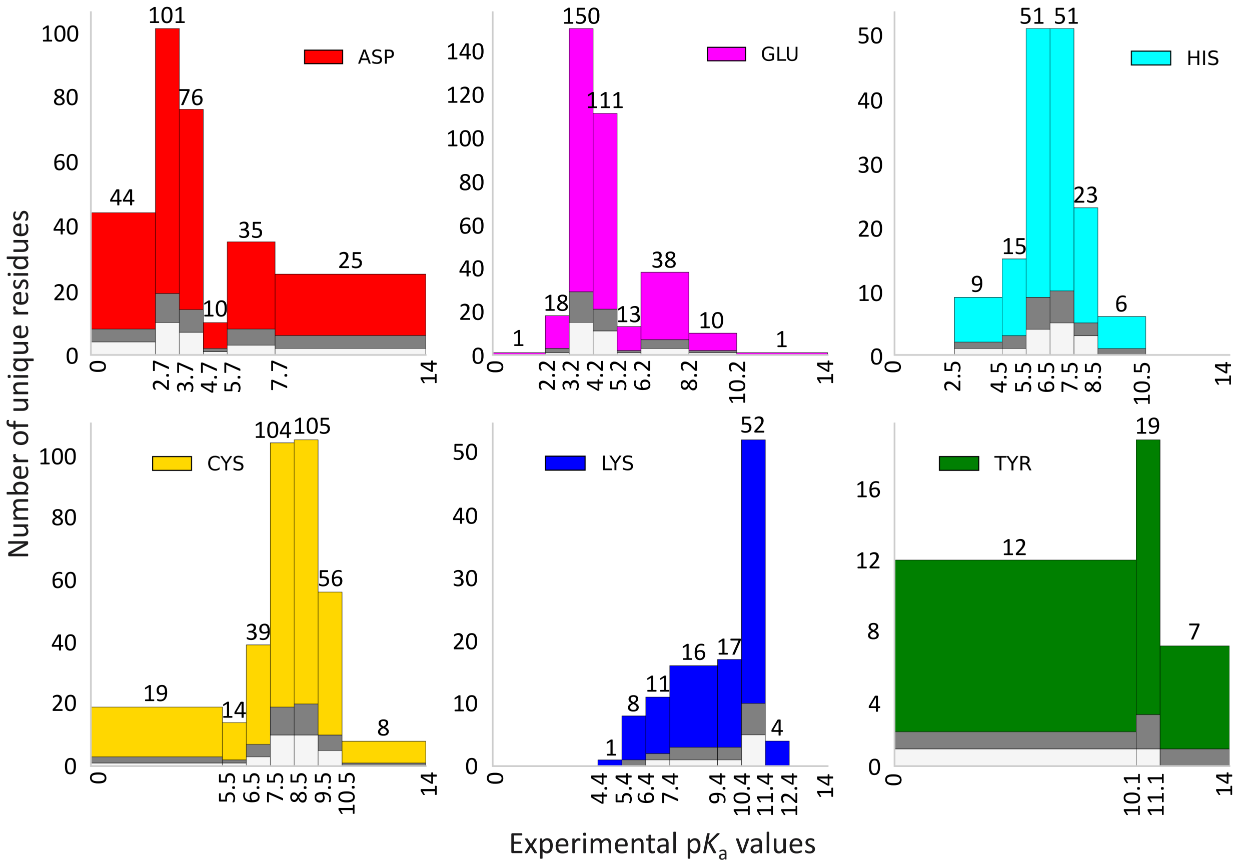


Figure S1: **Train-validation-test split.** The binning scheme used in this study for each residue type. The colored bins represent the overall dataset, grey bins represent the validation dataset, and the white bins represent the test dataset.

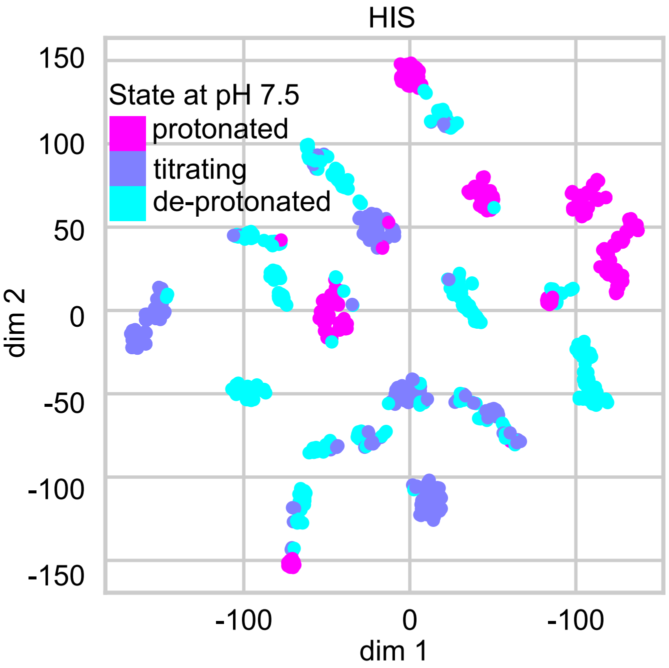


Figure S2: **t-distributed Stochastic Neighbor Embedding (t-SNE) visualization of the histidine data.** The numerical tree features of all histidines were extracted from the dataset with the AF2 augmentation. This 37 dimensional data was then mapped on two dimension using t-SNE. The color code indicates the protonation state at pH 7.0 based on the label experimental p$K_a$: protonated (p$K_a > 7.5$); deprotonated (p$K_a < 6.5$); titrating ( $6.5 \leq$ p$K_a \leq 7.5$).

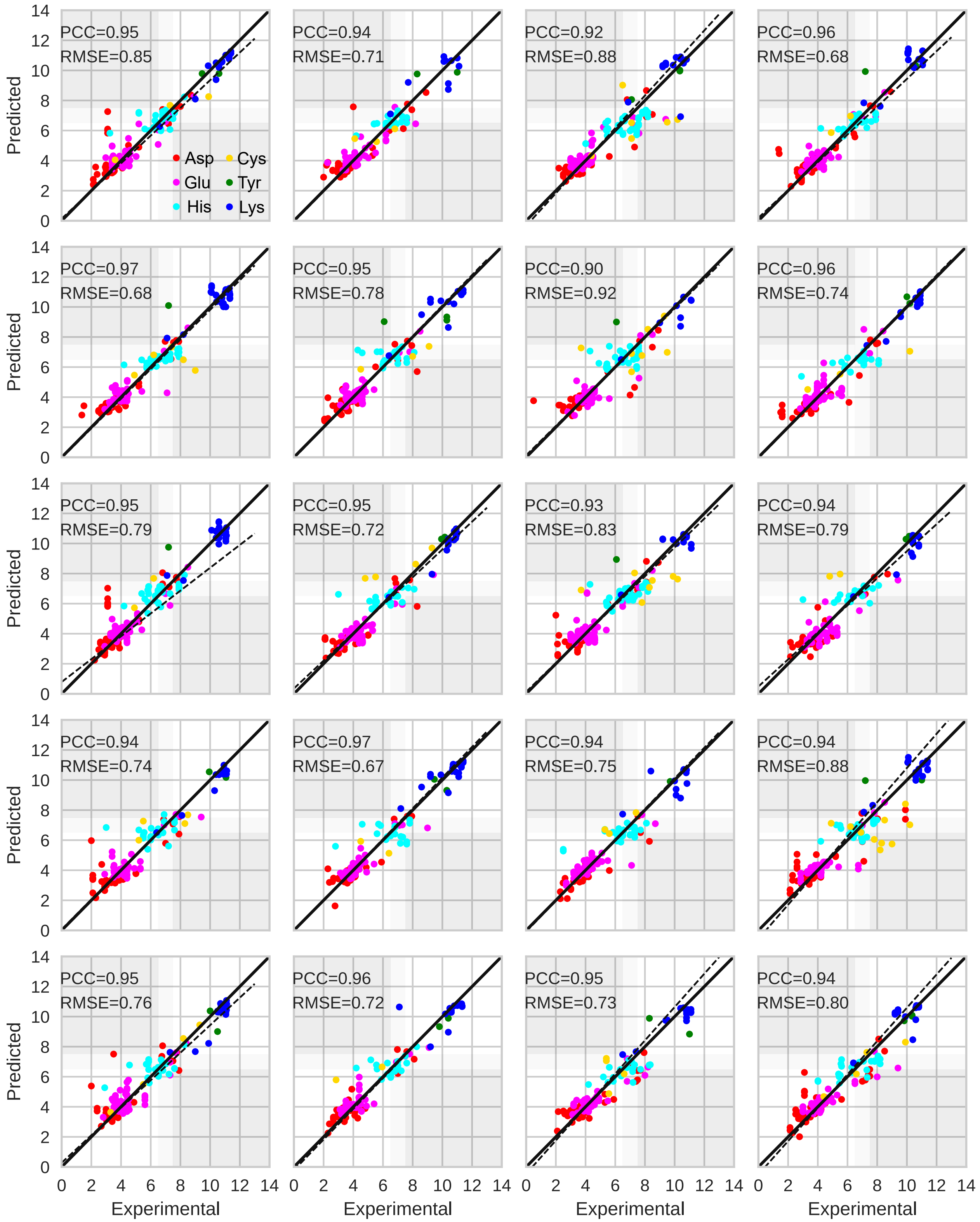


Figure S3: **Comparison of KaML-CBtree predicted and experimental p*K*$_a$'s in all 20 hold-out tests.** Predicted vs. experimental p*K*$_a$ values for all test splits. The solid line is the identity, the dashed line is a linear fit.

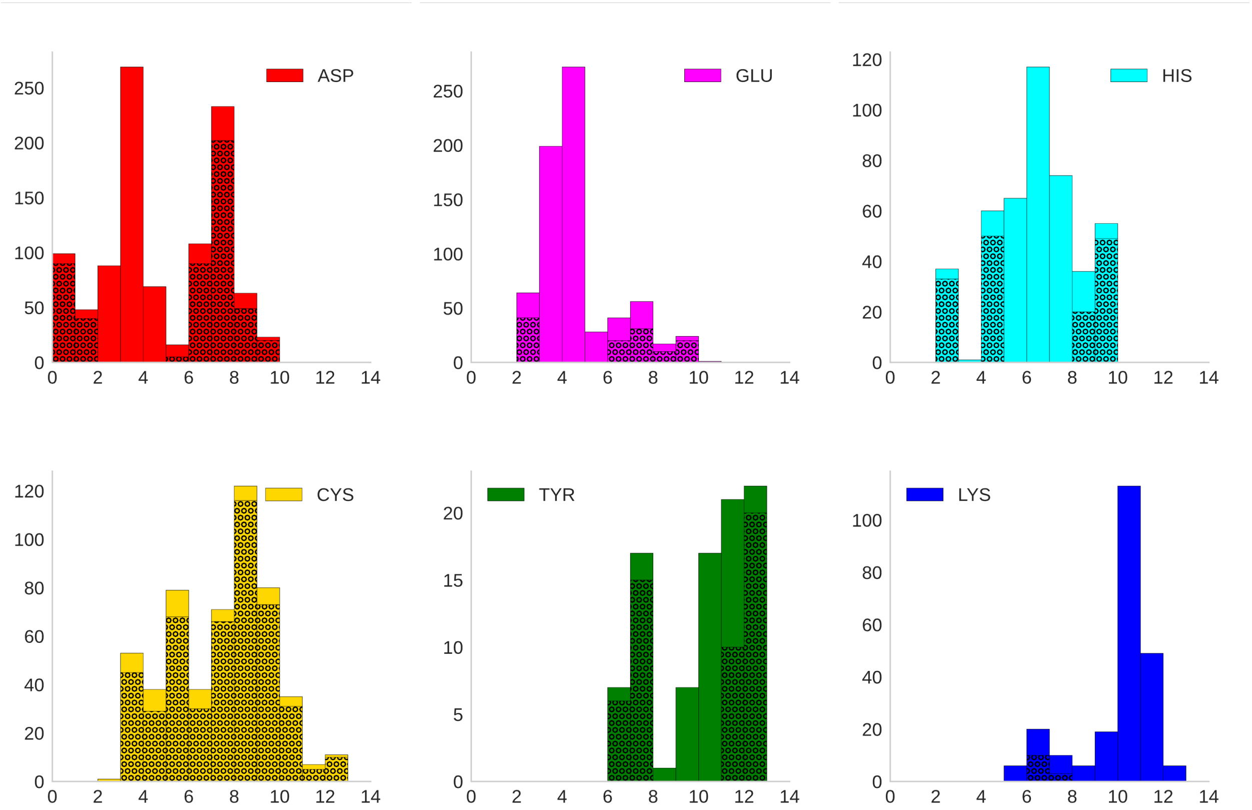


Figure S4: **Histograms of the p$K_a$ values of the training dataset after the AF2 augmentation.** Solid bars represent the total number of entries in the training dataset and shaded bars represent the augmentation using the AF2 structures.

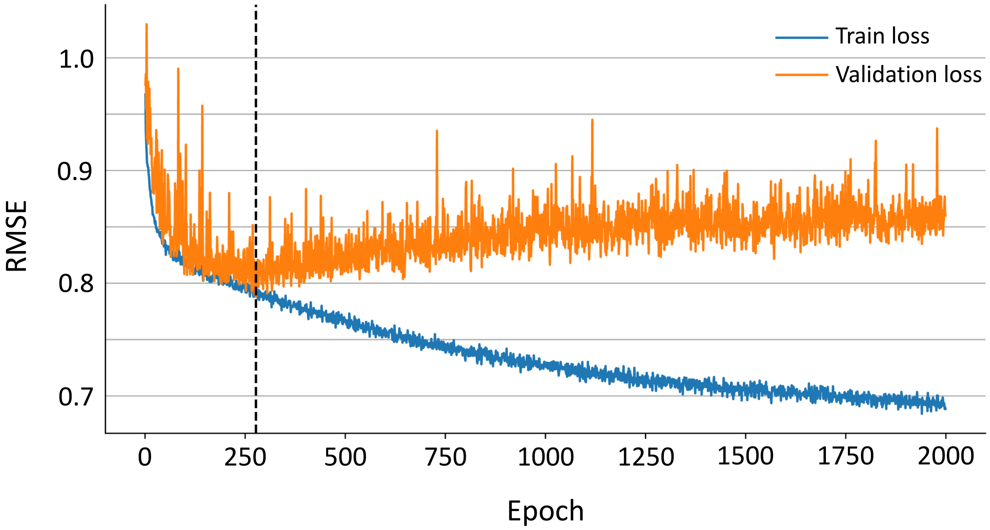


Figure S5: **Training and validation loss in the PHMD549 pre-training.** The dashed vertical line indicates the epoch where the model is saved. The training and validation RMSE is 0.79 and 0.79.

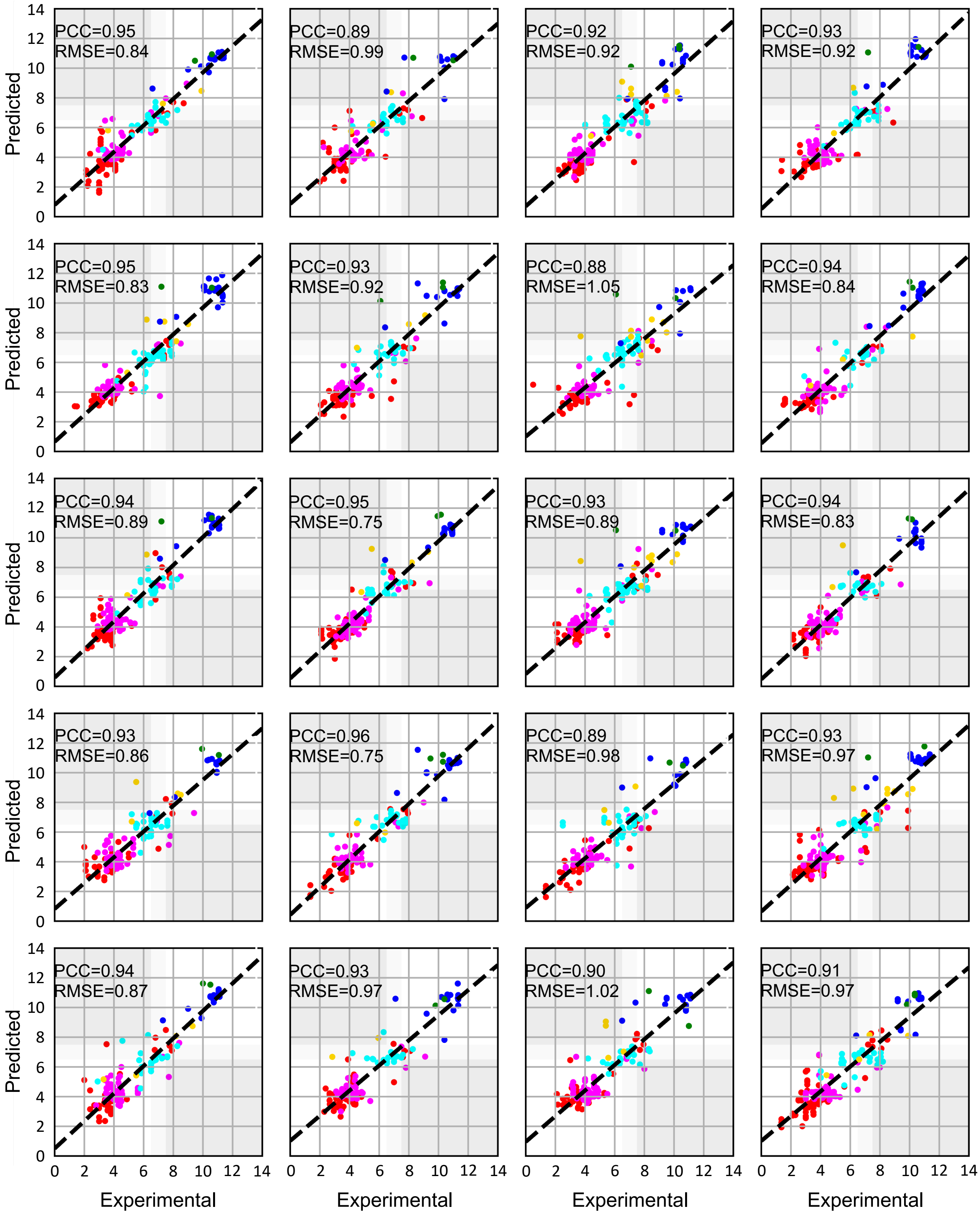


Figure S6: **Comparison of KaML-GAT predicted and experimental p$K_a$'s in all 20 hold-out tests.** Predicted vs. experimental p$K_a$ values for all test splits. The solid line is the identity, the dashed line is a linear fit.

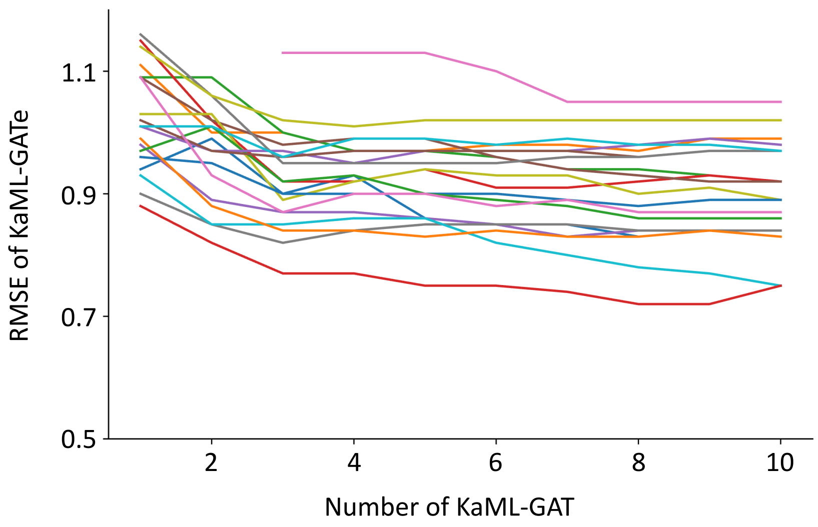


Figure S7: **The influence of the number of KaML-GAT models on the prediction performance of the ensemble KaML-GAT.** The x-axis is the number of KaML-GAT models in the ensemble KaML-GAT. The y-axis is the RMSEs of the ensemble KaML-GAT. The results for the 20 independent tests are shown in different colors.

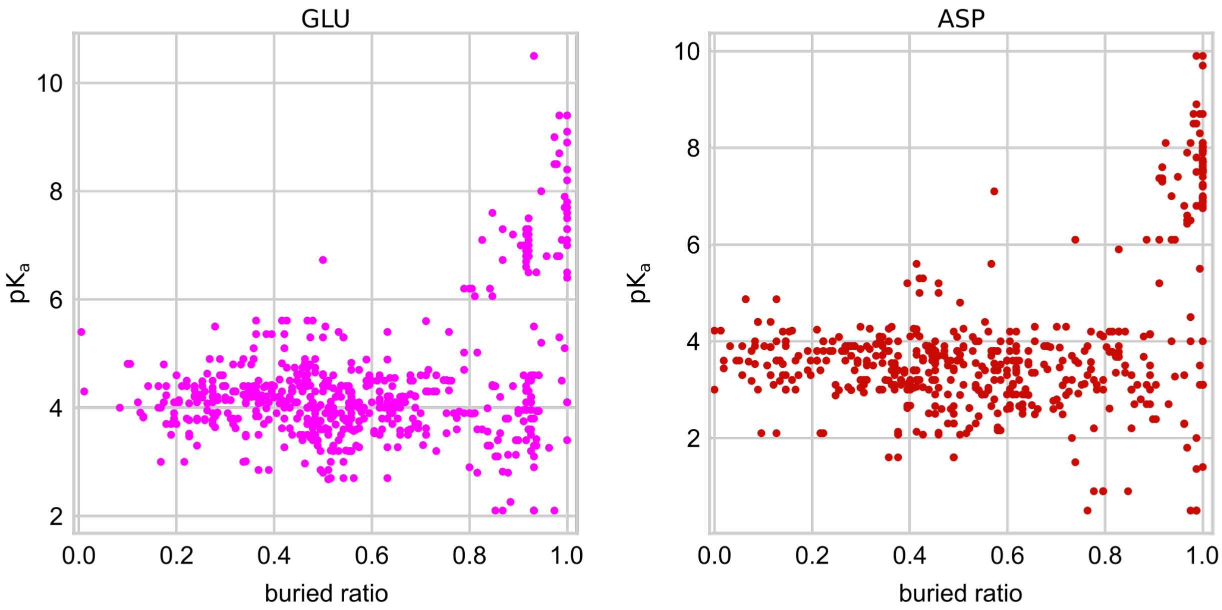


Figure S8: **Glu/Asp with large p$K_a$ values are deeply buried, but some deeply buried Asp/Glu show normal p$K_a$'s.** The experimentally determined p$K_a$ values vs. the calculated buried ratio.

(S18) The UniProt Consortium; Bateman, A.; Martin, M.-J.; Orchard, S.; Magrane, M.; Ahmad, S.; Alpi, E.; Bowler-Barnett, E. H.; Britto, R.; Bye-A-Jee, H.; Cukura, A.; Denny, P.; Dogan, T.; Ebenezer, T.; Fan, J.; Garmiri, P.; Da Costa Gonzales, L. J.; Hatton-Ellis, E.; Hussein, A.; Ignatchenko, A.; Insana, G.; Ishtiaq, R.; Joshi, V.; Jyothi, D.; Kandasaamy, S.; Lock, A.; Luciani, A.; Lugaric, M.; Luo, J.; Lussi, Y.; MacDougall, A.; Madeira, F.; Mahmoudy, M.; Mishra, A.; Moulang, K.; Nightingale, A.; Pundir, S.; Qi, G.; Raj, S.; Raposo, P.; Rice, D. L.; Saidi, R.; Santos, R.; Speretta, E.; Stephenson, J.; Totoo, P.; Turner, E.; Tyagi, N.; Vasudev, P.; Warner, K.; Watkins, X.; Zaru, R.; Zellner, H.; Bridge, A. J.; Aimo, L.; Argoud-Puy, G.; Auchincloss, A. H.; Axelsen, K. B.; Bansal, P.; Baratin, D.; Batista Neto, T. M.; Blatter, M.-C.; Bolleman, J. T.; Boutet, E.; Breuza, L.; Gil, B. C.; Casals-Casas, C.; Echioukh, K. C.; Coudert, E.; Cuche, B.; De Castro, E.; Estreicher, A.; Famiglietti, M. L.; Feuermann, M.; Gasteiger, E.; Gaudet, P.; Gehant, S.; Gerritsen, V.; Gos, A.; Gruaz, N.; Hulo, C.; Hyka-Nouspikel, N.; Jungo, F.; Kerhornou, A.; Le Mercier, P.; Lieberherr, D.; Masson, P.; Morgat, A.; Muthukrishnan, V.; Paesano, S.; Pedruzzi, I.; Pilbout, S.; Pourcel, L.; Poux, S.; Pozzato, M.; Pruess, M.; Redaschi, N.; Rivoire, C.; Sigrist, C. J. A.; Sonesson, K. et al. UniProt: The Universal Protein Knowledgebase in 2023. *Nucleic Acids Res.* **2023**, *51*, D523–D531.

(S19) Waterhouse, A.; Bertoni, M.; Bienert, S.; Studer, G.; Tauriello, G.; Gumienny, R.; Heer, F. T.; de Beer, T. A. P.; Rempfer, C.; Bordoli, L.; Lepore, R.; Schwede, T. SWISS-MODEL: Homology Modelling of Protein Structures and Complexes. *Nucleic Acids Res.* **2018**, *46*, W296–W303.

(S20) Jumper, J.; Evans, R.; Pritzel, A.; Green, T.; Figurnov, M.; Ronneberger, O.; Tunyasuvunakool, K.; Bates, R.; Žídek, A.; Potapenko, A.; Bridgland, A.; Meyer, C.; Kohl, S. A. A.; Ballard, A. J.; Cowie, A.; Romera-Paredes, B.; Nikolov, S.; Jain, R.; Adler, J.; Back, T.; Petersen, S.; Reiman, D.; Clancy, E.; Zielinski, M.; Steinegger, M.; Pacholska, M.; Berghammer, T.; Bodenstein, S.; Silver, D.; Vinyals, O.; Senior, A. W.; Kavukcuoglu, K.; Kohli, P.; Hassabis, D. Highly Accurate Protein Structure Prediction with AlphaFold. *Nature* **2021**, *596*, 583–589.

(S21) Mirdita, M.; Schütze, K.; Moriwaki, Y.; Heo, L.; Ovchinnikov, S.; Steinegger, M. ColabFold: Making Protein Folding Accessible to All. *Nat. Methods* **2022**, *19*, 679–682.

(S22) Steinegger, M.; Söding, J. MMseqs2 Enables Sensitive Protein Sequence Searching for the Analysis of Massive Data Sets. *Nat. Biotechnol.* **2017**, *35*, 1026–1028.

(S23) Pedregosa, F.; Varoquaux, G.; Gramfort, A.; Michel, V.; Thirion, B.; Grisel, O.; Blondel, M.; Prettenhofer, P.; Weiss, R.; Dubourg, V.; Vanderplas, J.; Passos, A.; Cournapeau, D.; Brucher, M.; Perrot, M.; Duchesnay, É. Scikit-Learn: Machine Learning in {P}ython. *J. Machine Learn. Res.* **2011**, *12*, 2825–2830.

(S24) Liu, R.; Clayton, J.; Shen, M.; Bhatnagar, S.; Shen, J. Machine Learning Models to Interrogate Proteome-Wide Covalent Ligandabilities Directed at Cysteines. *JACS Au* **2024**, *4*, 1374–1384.

(S25) Ali, M. PyCaret: An Open Source, Low-Code Machine Learning Library in Python. 2020.

(S26) Prokhorenkova, L.; Gusev, G.; Vorobev, A.; Dorogush, A. V.; Gulin, A. CatBoost: unbiased boosting with categorical features. 2019; https://arxiv.org/abs/1706.09516.

(S27) McElfresh, D.; Khandagale, S.; Valverde, J.; Feuer, B.; Hegde, C.; Ramakrishnan, G.; Goldblum, M.; White, C. When Do Neural Nets Outperform Boosted Trees on Tabular Data? 37th Conference on Neural Information Processing Systems. 2023.

(S28) Alcobaça, E.; Siqueira, F.; Rivolli, A.; Garcia, L. P. F.; Oliva, J. T.; de Carvalho, A. C. P. L. F. MFE: Towards reproducible meta-feature extraction. *Journal of Machine Learning Research* **2020**, *21*, 1–5.

(S29) Jamasb, A.; Viñas Torné, R.; Ma, E.; Du, Y.; Harris, C.; Huang, K.; Hall, D.; Lió, P.; Blundell, T. Graphein - a Python Library for Geometric Deep Learning and Network Analysis on Biomolecular Structures and Interaction Networks. Adv. Neural Inf. Process. Syst. 2022; pp 27153–27167.

(S30) Cai, Z.; Luo, F.; Wang, Y.; Li, E.; Huang, Y. Protein p$K_a$ Prediction with Machine Learning. *ACS Omega* **2021**, *6*, 34823–34831.

(S31) Cai, Z.; Liu, T.; Lin, Q.; He, J.; Lei, X.; Luo, F.; Huang, Y. Basis for Accurate Protein p$K_a$ Prediction with Machine Learning. *J. Chem. Inf. Model.* **2023**, *63*, 2936–2947.

(S32) Raschka, S. BioPandas: Working with Molecular Structures in Pandas DataFrames. *The Journal of Open Source Software* **2017**, *2*, 279.

(S33) Baber, J. C.; Hodgkin, E. E. Automatic Assignment of Chemical Connectivity to Organic Molecules in the Cambridge Structural Database. *J. Chem. Inf. Comput. Sci.* **1992**, *32*, 401–406.

(S34) MacKerell, A. D.; Bashford, D.; Bellott, M.; Dunbrack, R. L.; Evanseck, J. D.; Field, M. J.; Fischer, S.; Gao, J.; Guo, H.; Ha, S.; Joseph-McCarthy, D.; Kuchnir, L.; Kuczera, K.; Lau, F. T. K.; Mattos, C.; Michnick, S.; Ngo, T.; Nguyen, D. T.; Prodhom, B.; Reiher, W. E.; Roux, B.; Schlenkrich, M.; Smith, J. C.; Stote, R.; Straub, J.; Watanabe, M.; Wiórkiewicz-Kuczera, J.; Yin, D.; Karplus, M. All-Atom Empirical Potential for Molecular Modeling and Dynamics Studies of Proteins. *J. Phys. Chem. B* **1998**, *102*, 3586–3616.

(S35) Fey, M.; Lenssen, J. E. Fast Graph Representation Learning with PyTorch Geometric. International Conference on Learning Representations. 2019.

(S36) Veličković, P.; Cucurull, G.; Casanova, A.; Romero, A.; Liò, P.; Bengio, Y. Graph Attention Networks. International Conference on Learning Representations. 2018.

(S37) Polikar, R. Ensemble Based Systems in Decision Making. *IEEE Circuits Syst. Mag.* **2006**, *6*, 21–45.

(S38) Wallace, J. A.; Shen, J. K. Continuous Constant pH Molecular Dynamics in Explicit Solvent with pH-Based Replica Exchange. *J. Chem. Theory Comput.* **2011**, *7*, 2617–2629.

(S39) Harris, R. C.; Shen, J. GPU-Accelerated Implementation of Continuous Constant pH Molecular Dynamics in Amber: p$K_a$ Predictions with Single-pH Simulations. *J. Chem. Inf. Model.* **2019**, *59*, 4821–4832.

(S40) Glorot, X.; Bengio, Y. Understanding the Difficulty of Training Deep Feedforward Neural Networks. Proceed. Mach. Learn. Res. 2010.

(S41) Dayhoff II, G. W.; Uversky, V. N. Rapid Prediction and Analysis of Protein Intrinsic Disorder. *Protein Sci.* **2022**, *31*, e4496.

(S42) Kabsch, W.; Sander, C. Dictionary of Protein Secondary Structure: Pattern Recognition of Hydrogen-Bonded and Geomterical Features. *Biopolymers* **1983**, *22*, 2577–2637.

(S43) Reis, P. B. P. S.; Vila-Viçosa, D.; Rocchia, W.; Machuqueiro, M. PypKa: A Flexible Python Module for Poisson–Boltzmann-Based pKa Calculations. *J. Chem. Inf. Model.* **2020**, *60*, 4442–4448.
